# Multi-morph eco-evolutionary dynamics in structured populations

**DOI:** 10.1101/2021.07.08.451558

**Authors:** Sébastien Lion, Mike Boots, Akira Sasaki

## Abstract

Our understanding of the evolution of quantitative traits in nature is still limited by the challenge of including realistic trait distributions in the context of frequency-dependent selection and ecological feedbacks. We extend to class-structured populations a recently introduced “oligomorphic approximation” which bridges the gap between adaptive dynamics and quantitative genetics approaches and allows for the joint description of the dynamics of ecological variables and of the moments of multimodal trait distributions. Our theoretical framework allows us to analyse the dynamics of populations composed of several morphs and structured into distinct classes (e.g. age, size, habitats, infection status, species…). We also introduce a new approximation to simplify the eco-evolutionary dynamics using reproductive values. We illustrate the effectiveness of this approach by applying it to the important conceptual case of two-habitat migration-selection models. In particular, we show that our approach allows us to predict both the long-term evolutionary endpoints and the short-term transient dynamics of the eco-evolutionary process, including fast evolution regimes. We discuss the theoretical and practical implications of our results and sketch perspectives for future work.

---

Many experimental and empirical studies in evolutionary ecology aim at understanding how ecological processes affect how trait distributions change over time. This has motivated the development of quantitative genetics methods (QG) to analyse the dynamics of quantitative traits (Lande, 1979; Bulmer, 1992; Falconer, 1996; Walsh & Lynch, 2018). Following Lande (1976, 1979, 1982)’s seminal work, most quantitative genetics models assume unimodal trait distributions and frequency-independent selection, leaving aside the problem of how multimodal distributions can be generated by frequency-dependent disruptive selection. Under these assumptions, dynamical equations for the mean and higher moments of a trait distribution can be derived for tightly clustered trait distributions (Barton & Turelli, 1987; Turelli & Barton, 1990; Barton & Turelli, 1991). Assuming that trait distributions are narrowly localised around a single mean also allows one to incorporate frequency-dependence to some extent (Iwasa et al., 1991; Abrams et al., 1993), but typically models rely on the more classical assumption that trait distributions are and remain normally distributed. However, empirical evidence of skewed (Bonamour et al., 2017) or multimodal distributions highlight the need for an alternative approach.

A major limitation of current quantitative genetics theory is the reliance on simplified ecological scenarios that are not representative of the complexity of eco-evolutionary feedbacks in nature. This led to the development of adaptive dynamics theory (AD), which, under the assumption that evolution is limited by rare mutations, provides a mathematical framework to study the interplay between ecological and evolutionary processes (Metz et al., 1992; Dieckmann & Law, 1996; Metz et al., 1996; Geritz et al., 1998). Many authors have noted the similarities and subtle differences between AD and QG approaches under the assumptions of small mutational steps and narrow trait distributions respectively (Abrams et al., 1993; Abrams, 2001; Day, 2005; Lion, 2018c). However, there is a clear conceptual gap in the canonical approaches to AD and QG: while adaptive dynamics has been successful in taking into account environmental feedbacks and the emergence of polymorphism under frequency-dependent disruptive selection, it does so by assuming strong constraints on the mutation process and standing variation in the population. Recently, Sasaki & Dieckmann (2011) proposed an alternative “oligomorphic” approximation to bridge the gap between adaptive dynamics and quantitative genetics theory. The crux of the approach is to decompose a multimodal trait distribution into a sum of narrow unimodal morph distributions and to derive the dynamics of the frequency, mean trait value and variance of each morph. Suitable moment closure approximations at the morph level yield a closed dynamical system. As such, this framework can be seen as an extension of quantitative genetics theory to take into account eco-evolutionary feedbacks and polymorphic trait distributions.

The theoretical developments of Sasaki & Dieckmann (2011) rely on a number of additional assumptions, notably single-locus haploid genetics, large populations sizes, and unstructured populations. In this paper, we retain the first two assumptions but investigate how class structure affects the dynamics of quantitative trait distributions. As class structure is ubiquituous in biological populations, this is an important extension to Sasaki & Dieckmann (2011)’s theory, allowing us to apply the method to the majority of populations where individuals can be in distinct demographic, physiological or ecological states, such as different age groups, developmental stages, infection status, or habitats.

The paper is organised as follows. We first give a general decomposition of the trait distribution into different morphs in a population structured into distinct classes. We then show how, by assuming that each morph distribution is clustered around the morph mean, we can derive equations for the dynamics of class-specific morph frequencies, morph means, and morph variances using Taylor approximations of the vital rates describing between-classes transitions. We also apply recent theory on reproductive value (Lion, 2018a,b; Lion & Gandon, 2021) to simplify the morph dynamics at the population level. As in classical theory (Fisher, 1930; Taylor, 1990; Lehmann & Rousset, 2014; Gardner, 2015), a morph’s reproductive value is a measure of how well individuals of that morph in a given class transmit their genes to future generations, and we show that it can be used as a weight to calculate the net effect of selection on a given morph. Finally, we derive some simpler results for the important limit case of two-class models, and apply this general framework to three specific models describing the interplay between migration and selection in a population distributed over two habitats of distinct qualities coupled by migration. The first example revisits the local adaptation models analysed by Meszéna et al. (1997), Ronce & Kirkpatrick (2001), Débarre et al. (2013), and Mirrahimi & Gandon (2020), but our approach allows us to express these previous results in terms of the reproductive values of each habitat and to take into account habitat-specific mutation. The second example is a two-habitat extension of the resource competition model analysed by Sasaki & Dieckmann (2011) and highlights how our framework can shed light on frequency-dependence and disruptive selection. The third example is a resource-consumer model and is used to show that our approach allows us to analyse transient eco-evolutionary dynamics fuelled by non-neligible standing variation at the population level. Together these examples illustrate the potential of the approach in a wide range of fundamental eco-evolutionary scenarios.

## 1 Densities and trait distributions

We consider a population of individuals characterised by a continuous phenotypic trait. The total density of individuals with trait value *z* at time *t* is *n*(*z*, *t*) and, for simplicity, we denote the total density of individuals as *n*(*t*) = *∫ n*(*z*, *t*)d*z* (with a slight abuse of notation). We further assume that the population is structured into *K* discrete classes, which can for instance represent different age groups, developmental stages, or habitats. The density of individuals with trait value *z* in class *k* at time *t* is *n^k^*(*z*, *t*). Similarly, we write *n^k^*(*t*) = *∫ n^k^*(*z*, *t*)d*z* for the total density of individuals in class *k* at *t*. See Table 1 for a description of the main notations in the paper.

**Table 1:**
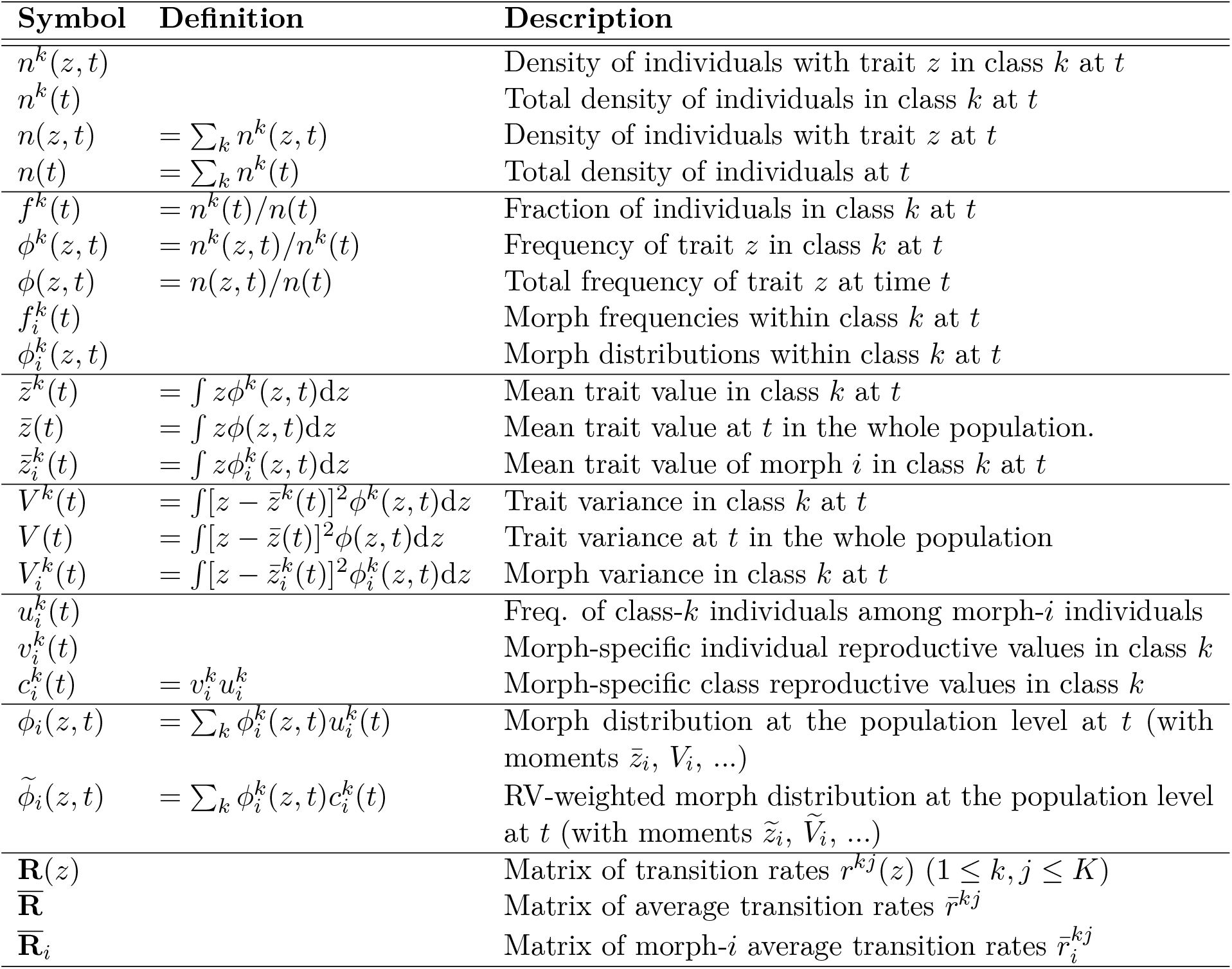
Definition of mathematical symbols used in the text

| Symbol | Definition | Description |
| --- | --- | --- |
| $n^k(z, t)$ | | Density of individuals with trait $z$ in class $k$ at $t$ |
| $n^k(t)$ | | Total density of individuals in class $k$ at $t$ |
| $n(z, t)$ | $= \sum_k n^k(z, t)$ | Density of individuals with trait $z$ at $t$ |
| $n(t)$ | $= \sum_k n^k(t)$ | Total density of individuals at $t$ |
| $f^k(t)$ | $= n^k(t)/n(t)$ | Fraction of individuals in class $k$ at $t$ |
| $\phi^k(z, t)$ | $= n^k(z, t)/n^k(t)$ | Frequency of trait $z$ in class $k$ at $t$ |
| $\phi(z, t)$ | $= n(z, t)/n(t)$ | Total frequency of trait $z$ at time $t$ |
| $f_i^k(t)$ | | Morph frequencies within class $k$ at $t$ |
| $\phi_i^k(z, t)$ | | Morph distributions within class $k$ at $t$ |
| $\bar{z}^k(t)$ | $= \int z \phi^k(z, t) dz$ | Mean trait value in class $k$ at $t$ |
| $\bar{z}(t)$ | $= \int z \phi(z, t) dz$ | Mean trait value at $t$ in the whole population. |
| $\bar{z}_i^k(t)$ | $= \int z \phi_i^k(z, t) dz$ | Mean trait value of morph $i$ in class $k$ at $t$ |
| $V^k(t)$ | $= \int [z - \bar{z}^k(t)]^2 \phi^k(z, t) dz$ | Trait variance in class $k$ at $t$ |
| $V(t)$ | $= \int [z - \bar{z}(t)]^2 \phi(z, t) dz$ | Trait variance at $t$ in the whole population |
| $V_i^k(t)$ | $= \int [z - \bar{z}_i^k(t)]^2 \phi_i^k(z, t) dz$ | Morph variance in class $k$ at $t$ |
| $u_i^k(t)$ | | Freq. of class-$k$ individuals among morph-$i$ individuals |
| $v_i^k(t)$ | | Morph-specific individual reproductive values in class $k$ |
| $c_i^k(t)$ | $= v_i^k u_i^k$ | Morph-specific class reproductive values in class $k$ |
| $\phi_i(z, t)$ | $= \sum_k \phi_i^k(z, t) u_i^k(t)$ | Morph distribution at the population level at $t$ (with moments $\bar{z}_i, V_i, \dots$) |
| $\tilde{\phi}_i(z, t)$ | $= \sum_k \phi_i^k(z, t) c_i^k(t)$ | RV-weighted morph distribution at the population level at $t$ (with moments $\tilde{z}_i, \tilde{V}_i, \dots$) |
| $\mathbf{R}(z)$ | | Matrix of transition rates $r^{kj}(z)$ ($1 \leq k, j \leq K$) |
| $\bar{\mathbf{R}}$ | | Matrix of average transition rates $\bar{r}^{kj}$ |
| $\bar{\mathbf{R}}_i$ | | Matrix of morph-$i$ average transition rates $\bar{r}_i^{kj}$ |

### 1.1 Full distributions

The within- and across-class densities represent the raw statistics of the model. They can be used to define some useful distributions to analyse the eco-evolutionary dynamics of the population. At the ecological level, the **class distribution** can be defined as

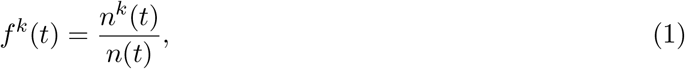

which represents the fraction of individuals that are in class *k* at time *t*. Note that ∑_*k*_ *f^k^*(*t*) = 1, where the summation is over all classes, i.e. 1 ≤ *k* ≤ *K* (for simplicity, all summation limits will be implicit in this article).

At an evolutionary level, two **trait distributions** can be defined. The within-class trait distribution is

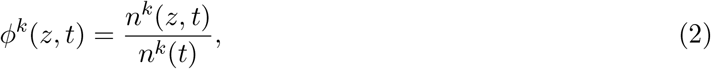

which is the frequency of individuals with trait *z* in class *k* at time *t*. Averaging over classes yields the across-class trait distribution

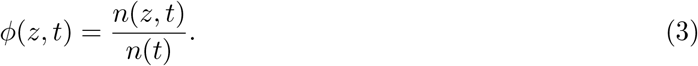

It is easy to check that, as expected, *∫ ϕ^k^*(*z*, *t*)d*z* = *∫ ϕ*(*z*, *t*)d*z* = 1. Note that the class and trait distributions are linked through the relationship *ϕ*(*z*, *t*) = ∑_*k*_*ϕ^k^*(*z*, *t*)*f^k^*(*t*).

### 1.2 Multi-morph decomposition

Up to now, we have made no assumption on the trait distribution in the population. With the notations defined so far, it is straightforward to produce a continuous-trait version of the Price equations derived in Lion (2018a), but our aim here is slightly different, because we want to make specific predictions on the dynamics of multimodal distributions. Following Sasaki & Dieckmann (2011), we therefore assume that the trait distribution can be decomposed into *M* morphs. Specifically, we use the term ‘morph’ to describe a cluster of continuous variants around a phenotypic mean trait. We allow morphs to have distinct distributions in different classes, and write the within-class trait distributions as a mixture of morph distributions,

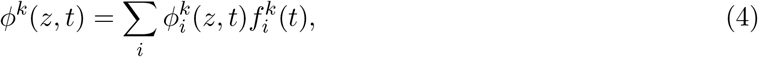

where 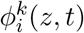 is the distribution of morph *i* in class *k* at time *t*, and 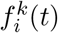 is the frequency of morph *i* in class *k*. Note that 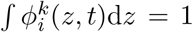 and 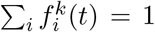, where the summation is implicitly over all morphs (i.e. 1 ≤ *i* ≤ *M*). Equation (4) is a class-specific version of equation (5) in Sasaki & Dieckmann (2011). Biologically it means that the full distribution *ϕ^k^*(*z*, *t*) can be decomposed into a sum of morph distributions, 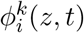, each weighted by the morph frequency, 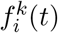. Figure 1 gives a graphical illustration of the multi-morph decomposition using simulation results from a two-class, two-morph example (e.g. Example 1 below).

**Figure 1:**
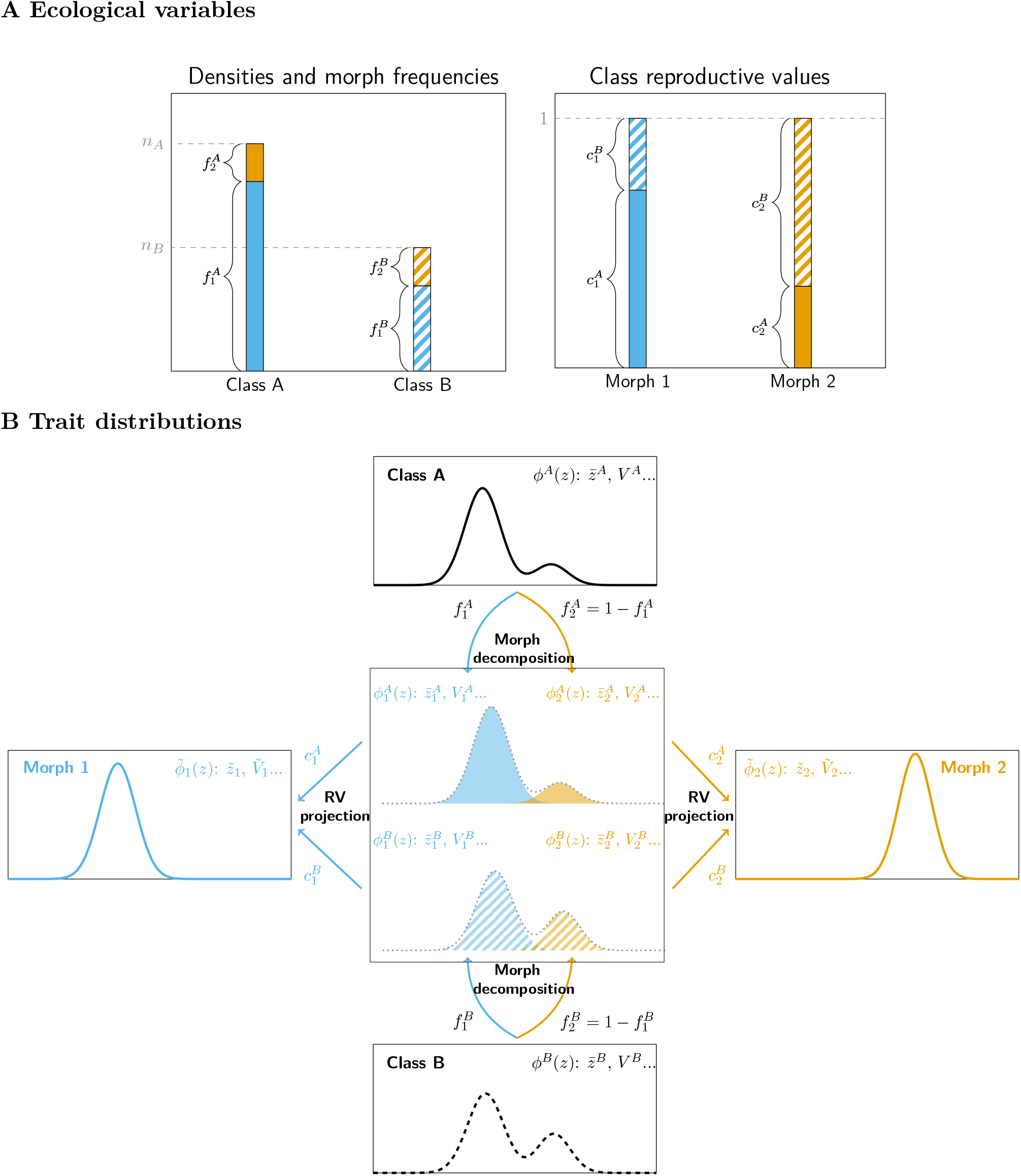
Summary of the notations and approach using a two-class, two-morph example. Panel (a) shows the fast variables, which change on the ecological time scale. These are the densities of individuals in each class, *n^A^*(*t*) and *n^B^*(*t*), the morph frequencies in each class, 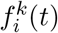, and the morph-specific class reproductive values, 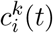. In the simulation snapshot used to plot these graphs, morph 1 is relatively more abundant within class B 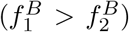, but has a lower class reproductive value 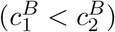. Panel (b) shows the trait distributions, which change on the slow, evolutionary time scales. The trait distribution in class A, *ϕ^A^*(*z*, *t*), can be decomposed into a mixture of class-specific morph distributions, 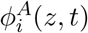, weighted by the class-specific morph frequencies 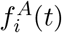. Note that, to better illustrate the decomposition, the shaded areas represent 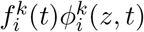, and not the distributions 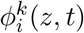. This multi-morph decomposition can also be applied to class *B*. On the slow time scale, the relevant aggregate distributions at the morph level are the RV-weighted morph distributions 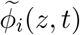. (Note that the graphs use data from a numerical simulation of Example 1.)

Intuitively, it makes sense to associate the distribution 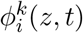 to one “peak” of a multimodal distribution, but it is important to note that the decomposition (4) is also valid if the morph distributions overlap or are similar. We can for instance start with two very similar morphs and study how disruptive selection causes the unimodal distribution *ϕ^k^*(*z*, *t*) to split into two peaks as the morph means move away from each other. Our theoretical framework therefore does not require the distance between morph means to be large. Similarly, our analysis does not require a morph to have the same distribution or the same frequency in all classes, although it will often make biological sense to consider scenarios where the trait distributions in the different classes look similar (without being identical). Finally, while the number of morphs can be arbitrarily chosen, it makes sense to use biological intuition to guide this choice (for instance, two morphs for a two-resource model; see the discussion for a more formal argument).

### 1.3 Morph moments

From the distributions *ϕ^k^*(*z*, *t*), we can calculate class-specific moments, such as 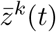, the mean trait value in class *k* at time *t*, and *V^k^*(*t*), the trait variance in class *k* at time *t*. Similarly, morph-specific moments can be calculated from the distributions 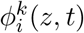. For instance, the mean trait value of morph *i* in class *k* at *t* is 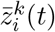, and the trait variance of morph *i* in class *k* at *t* is 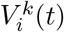.

If higher-order moments are negligible or can be approximated using moment closure approximations at the morph level, a morph can then be characterised by its relative abundance (e.g. its frequency in each class, 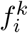), its position (e.g. the class-specific morph mean 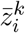), and its width (e.g. the class-specific morph standard deviation 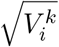). Equation (4) allows us to make connections between population-level moments and morph-specific moments. See Table 1 for explicit definitions of the population-level and morph-specific moments, as well as figure 1 for a graphical summary of the notations.

### 1.4 Notational conventions

To simplify the notations, a number of conventions will be used throughout the paper. First, classes will be identified by superscripts and morphs by subscripts. For classes, we use the superscripts *j* or *k*, so that an implicit summation over *k* means that *k* takes values between 1 and *K*. For morphs, we use the subscript *i*, which thus takes values between 1 and *M*. The symbol *ℓ* will be used either for classes or morphs, when needed. Second, whenever it is clear from the context, we shall drop the dependency on time, writing e.g. *f^k^* instead of *f^k^*(*t*).

## 2 Dynamics and separation of time scales

Having defined the statistics we need to describe the state of the population at a given time, we now turn to their dynamics. Figure 1 illustrates how a multimodal trait distribution can be decomposed into a mixture of unimodal morph distributions. By tracking the dynamics of these morph distributions, we can understand how the various peaks of the multimodal distribution move and change over time. To derive these dynamics, we first specify the rates associated with the different events of the life cycle, then we calculate an approximation of these rates under the assumption that the morph distributions are sufficiently narrow.

### 2.1 Vital rates

At a general level, the vital rates are defined by functions *r^jk^*(*z*, *E*(*t*)), which give the rate of production of individuals in class *j* by an individual in class *k* with trait *z* at time *t*. The variable *E*(*t*) represents the environmental feedback, which collects all ecological variables needed to calculate the reproduction and survival of individuals (Metz et al., 1992; Mylius & Diekmann, 1995; Metz et al., 2008; Lion, 2018c). For clonally reproducing organisms, this is sufficient to calculate the dynamics of the density *n^j^*(*z*, *t*), as follows

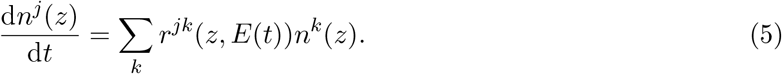

Equation (9) describes the interplay of selection and ecological dynamics under the assumption of perfectly faithful reproduction (that is individuals with trait *z* give birth to individuals with trait *z*). In our initial presentation, we find it easier to omit the mutation process, but we will consider the effect of mutation at a later stage. In addition, we do not explicitly model class-specific traits in this paper (but see the discussion for how this can be taken into account).

In the remainder of the manuscript, all operations on the vital rates *r^jk^*(*z*, *E*(*t*)) will be partial derivatives or integration with respect to the first argument. Hence, we shall drop the dependency on environmental feedback and write simply *r^jk^*(*z*) (and **R**(*z*) for the matrix of vital rates). However, it must be kept in mind that this notation does not imply density-independent or frequency-independent selection. As shown in the “Applications” section, our formalism can be readily applied to scenarios where the vital rates depend on the density of conspecifics or other species, on the trait distribution, or on other biotic or abiotic ecological variables (see e.g. Sasaki & Dieckmann (2011) and Lion (2018c)).

### 2.2 Small morph variance approximation

The crux of the oligomorphic approximation of Sasaki & Dieckmann (2011) is to assume that the morph distributions are tightly clustered around their mean, that is the standard deviation of the morph distribution is proportional to a small parameter *ε*. In a class-structured model, this means that the quantity 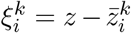 is small, and we write 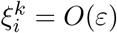. A simple Taylor expansion of the vital rates *r^jk^*(*z*) around the within-class morph mean 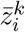 yields

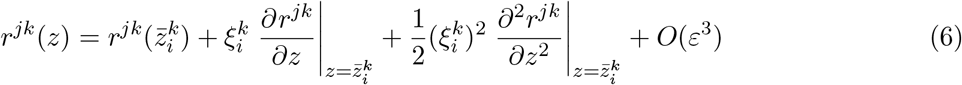

Integrating over the distribution 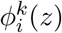 yields an approximation for the average vital rates of morph *i*, in terms of the morph-specific mean and variances 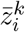 and 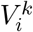. We have

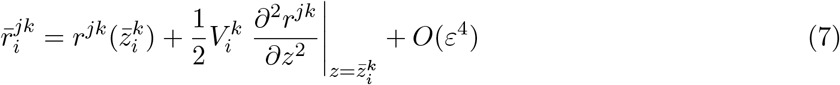

Similarly, averaging *r^jk^*(*z*) over the distribution *ϕ^k^*(*z*) yields the average vital rates 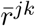, which can be decomposed in terms of morph averages as 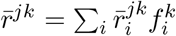. Note that, as in Sasaki & Dieckmann (2011), we assume that the morph distributions are symmetric around their mean throughout this paper (which is why the remainder in equation (7) is of order *ε*^4^).

### 2.3 Separation of time scales

Because we assume that within-morph variation is small, the oligomorphic approximation gives rise to a separation of time scales between ecological and evolutionary variables. Specifically, the dynamics of the densities *n^k^*, class frequencies *f^k^* and morph frequencies 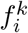 are all *O*(1) so that these can be treated as fast variables. On the other hand, the dynamics of morph means and variances will tend to vanish as *ε* tends towards 0, so the morph moments change on slower time scales. It is important to realise that this does not mean that there is no feedback between ecology and evolution, and in fact this approximation can be used to study situations where rapid evolution is fuelled by a large standing variance at the population level (for instance if we have two morphs with very different mean trait values), while assuming that the standing variation *in each morph* remains small. This will be explored in our Example 3 below.

Thus, the resulting coupled dynamics of densities, morph frequencies, morph means and morph variances take the form of a fast-slow dynamical system (see Rinaldi & Scheffer (2000) for an ecologically oriented overview). There are two ways to intepret this system. First, it can be viewed as an approximation of the full eco-evolutionary dynamics, and can either be numerically explored or used to gain analytical insight on the transient and long-term dynamics of the ecological and evolutionary variables, as is typically carried out in quantitative genetics approaches. Second, it can be analysed using quasi-equilibrium approximations, following the typical practice in evolutionary invasion analyses such as adaptive dynamics. This dictates that one first derives the equilibrium of the ecological variables and morph frequencies, for fixed values of the morph means and variances, then uses this information to calculate the dynamics of the morph means, the evolutionary singularities and the dynamics of morph variances near these singularities.

In the following, we first derive the equations of the fast variables (the densities and morph frequencies), then those of the slow variables (the morph means and variances). This allows us to clarify the connections with previous approaches. For instance, the equations of the morph frequencies are reminiscent of those governing allele frequency change in classical population genetics models, except that in our approach the means and variances are not fixed. Similarly, the dynamics of the morph means are reminiscent of those of classical quantitative genetics, but we go one step further by deriving the dynamics of morph variances instead of assuming that they are fixed. In this way, our approach relaxes some key assumptions of classical population and quantitative genetics, while providing a more dynamical perspective than classical invasion analyses.

## 3 Dynamics of ecological variables and morph frequencies

In this section, we derive the dynamics of the fast variables, which are the class densities *n^k^*(*t*) and class-specific morph frequencies 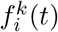. We also introduce the idea of weighting each class by its reproductive value in order to calculate the net effect of selection on the change of frequency of a given morph.

### 3.1 Dynamics of densities

Collecting all the class densities *n^k^*(*t*) in a vector **n**, we can write

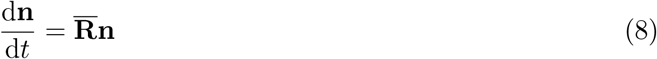

where 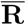 is the matrix of average vital rates 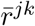. Using (7), we have 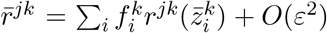, so the dynamics of class densities only depend, to zeroth order, on the morph frequencies and means. More explicitly, we have

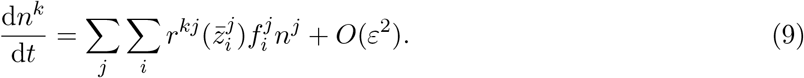

Similarly, the vector of class frequencies **f** = **n**/*n* has the following dynamics (Lion, 2018a)

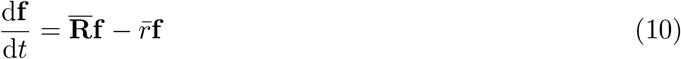

where 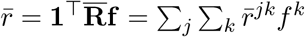 is the average growth rate of the total population. Again, expansion (7) shows that the dynamics of class frequencies is *O*(1) and solely determined by the morph frequencies and morph means.

### 3.2 Dynamics of morph frequencies

In Appendix A, we show that the dynamics of the within-class morph frequencies 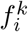 can be written as

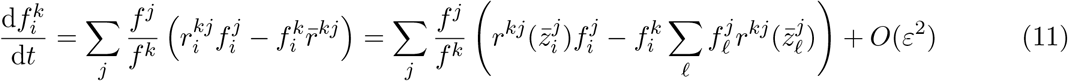

which shows that the morph frequencies 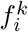 also have fast dynamics that depend only on the morph positions and frequencies. Equation (11) is the class-structured extension of the first line of equation (17) in Sasaki & Dieckmann (2011) and is a class-structured version of the replicator equation (Crow & Kimura, 1970; Ewens, 2004).

Later, we shall see that it is also useful to introduce the total frequency of morph *i*, 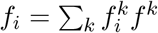, and the vector **u**_*i*_ collecting the morph frequencies 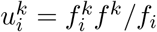, which gives the fraction of morph-*i* individuals which are in class *k*. The dynamics of **u**_*i*_ is then

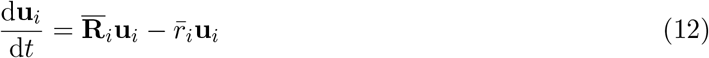

where 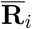 is the matrix of morph-specific average rates, 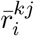, and 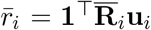 is the average growth rate of morph *i*. Hence, equation (12) is the morph-specific version of equation (10). Note the difference between the two morph frequencies 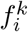 and 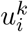. The frequency 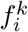 gives the fraction of morph-*i* individuals among all class-*k* individuals, while the frequency 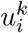 is the fraction of class-*k* individuals among all morph-*i* individuals.

### 3.3 Dynamics of morph reproductive values

As in Lion (2018a), equation (12) has a “companion” equation (or adjoint equation, in mathematical terms), which gives the dynamics of the vector of individual reproductive values for morph *i*, **v**_*i*_

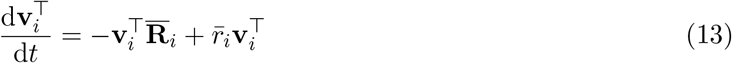

The reproductive value 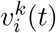 measures the relative contribution to the future of a morph-*i* individual in class *k* at time *t*, and therefore gives an instantaneous measure of the relative quality of class *k* from the point of view of morph *i*. Note that **v**_*i*_ and **u**_*i*_ are co-normalised such that 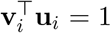. This co-normalisation condition means that the average quality of a morph-*i* individual is 1 at all times (Lion & Gandon, 2021). Later in the paper, we will show that we can weight individuals in different classes by their reproductive values in order to calculate the net effect of selection on morph *i*, which will allow us to reduce the dimension of the eco-evolutionary dynamics.

## 4 Dynamics of morph means and variances

On the fast time scale where morph means and variances do not change much, (9) and (11) are sufficient to describe the eco-evolutionary dynamics of the population. We now look at the slower time scales corresponding to changes in the morph means 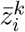 and morph variances 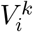.

### 4.1 Dynamics of morph means

In Appendix B, we show that the dynamics of morph means take the form of a class-structured Price equation (similar to those derived in Lion (2018a,b)). Using expansion (6), and assuming that the morph distributions are and remain symmetric, we obtain

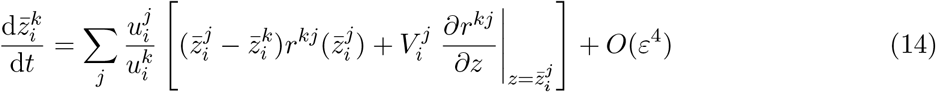

The first term between brackets in equation (14) represents the effect of demographic transitions between classes, and can be viewed as a “migration” term. In the absence of other processes, the phenotypic differentiation 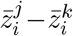 will tend to be consumed by demographic transitions from classes *j* to class *k*, which occur at rates 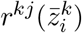. However, selection itself can generate phenotypic differentiation. The second term between brackets in equation (14) corresponds to the effect of directional selection on morph *i* within class *j*, and depends on the variance 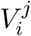 of morph *i* in class *j* and on the marginal effect of the trait on the vital rates, evaluated at the morph mean in class *j*. Finally, the ratio 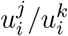 gives the relative abundance of class *j* and *k* in the population of morph-*i* individuals and is used as a weight to obtain the net change of the morph mean in class *k*, so that classes with a low frequency in the population do not contribute much. Note that, in the absence of class structure, the first term between brackets vanishes and we recover equation (25) in Sasaki & Dieckmann (2011).

It may not be immediately obvious that 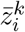 is a slow variable. Indeed, although 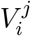 is *O*(*ε*^2^) by assumption, the first term between brackets is not necessarily small. For many biologically realistic applications, however, we can further assume that the morph means in the different classes are not too different, and more precisely that the phenotypic differentiation 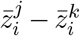 is *O*(*ε*), in which case the 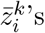 will change on a slower time scale compared to the morph frequencies 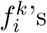, and the class densities *n^k^*. In a quasi-equilibrium approximation, this means that, while the 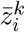 change slowly, the fast variables immediately track this change so that the right-hand sides of equations (9)–(11), which all explicitly depend on the morph means 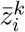, can be set to zero. More generally, we can use a perturbation expansion to show that the 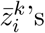 have a fast component (corresponding to the homogenisation of morph means due to between-class demographic transitions) and a slow component (corresponding to the effect of selection), so that we can relax the assumption that the differentiations 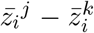 are initially small (Online Appendix S.3).

### 4.2 Dynamics of morph variances

A classical quantitative genetics approach would typically focus on equations (14) under the assumption of constant variances. However, this is not sufficient to understand how disruptive selection may shape multi-morph trait distributions, and for this we need to turn to the dynamics of the morph variances.

In Appendix B, we show that the dynamics of the class-specific morph variances can be written as:

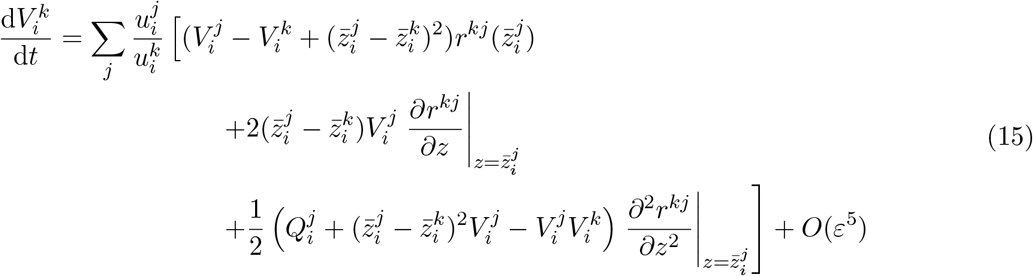

The term on the first line corresponds to the effect of demographic transitions between classes on variance. Basically it tells us that, even in the absence of selection, changes in the morph variance in class *k* can be observed if the morph distributions are different across classes (e.g. if the morph mean and variance in class *j* differ from class *k*). The term on the second line represents the effect of directional selection on the morph variance in class *k*, which will be greater when there is substantial phenotypic differentiation between the focal class and the other classes. Finally, the term on the third line represents the effect of disruptive selection on the morph variance, and depends on the means, variances, and fourth central moments of the morph distributions in class *j*, 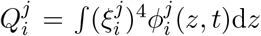. Note that, in the absence of class structure, the first two lines vanish and we recover equation (33) in Sasaki & Dieckmann (2011).

If we assume, as in the previous section, that 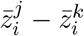 is *O*(*ε*), then the terms on the first, second and third lines are respectively *O*(*ε*^2^), *O*(*ε*^3^) and *O*(*ε*^4^), and it is immediately clear that the morph variances change on a slow time scale compared to the morph densities and frequencies. A more general argument can be obtained using the perturbation expansion approach detailed in Online Appendix S.3, as for the dynamics of the means.

### 4.3 Moment closure

As typical of moment methods, the dynamics of morph variances (15) depend on higher-order moments, notably the fourth central moments 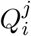. Hence, we need to close the system using a suitable moment closure approximation. For unstructured populations, Sasaki & Dieckmann (2011) studied two moment closure approximations, namely the Gaussian approximation and the house-of-cards approximation. In this paper, we focus on the Gaussian approximation, and therefore assume that the morph distributions 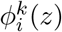 are normal. Biologically, this means that the total distribution of the trait can be viewed as a weighted sum of normal distributions. Mathematically, this entails that 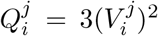, which is sufficient to close the system. Importantly, the assumption of normal morph distributions only matters when calculating the dynamics of variances.

### 4.4 Adding mutation

So far, we have analysed the effect of selection on the dynamics of densities, morph frequencies, morph means and variances, in the absence of mutation. There are several ways to incorporate mutation in our framework, but the simplest is to assume that mutation is independent of birth events and is unbiased at the morph level. With these assumptions, we show in Appendix E that the only effect of mutation is to increase morph variance by a term 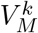, which is the mutational variance in class *k* (that is the product of the mutation rate *μ^k^* times the variance of the mutation kernel in class *k*). Thus, any depletion of morph variance due to stabilising selection will tend to be regenerated by mutation.

## 5 Morph-level dynamics: projection on RV space

The above analysis allows us to derive dynamical equations for the ecological densities, morph frequencies, morph means and morph variances, but this results in a high-dimensional system. In this section, we show that we can reduce the dimension of this system by tracking the moments of average morph distributions, where the average is obtained by weighting each class by both its quality and quantity. There are several possible weightings, but the most convenient is to use reproductive values (RV) as weights, as this choice leads to simpler equations and ensures that we capture the net effect of selection on a given morph (Fisher, 1930; Taylor, 1990; Lehmann & Rousset, 2014; Gardner, 2015; Grafen, 2015; Lion, 2018a).

### 5.1 RV-weighted trait distributions

We use the reproductive values we introduced in section 3.3 to define a weighted trait distribution (Lion, 2018a)

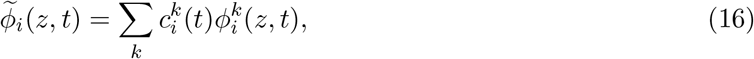

where 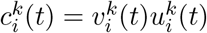 is the class reproductive value of morph *i* in class *k* at time *t*. Equation (16) weighs the class-specific morph distribution by both the quantity 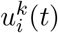 and quality 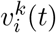 of class-*k* individuals within the morph-*i* subpopulation. As intuitively expected, the quality and quantity of a class may change over time, so these are dynamical variables (Lion, 2018a; Lion & Gandon, 2021). Figure 1 gives a graphical illustration of the process by which class- and morph-specific distributions can be aggregated to obtain morph-specific RV-weighted distributions.

Biologically, equation (16) gives us an appropriate metric to measure the average effect, across all classes, of selection acting on a given morph. By accounting for the relative qualities of each class, it is possible to get rid of any spurious effect due to intrinsic demographic differences between classes (e.g. when one class has a systematically higher productivity than the others, irrespective of the individuals’ genotypes). Any observed change in the RV-weighted trait distribution will therefore be the result of selection only.

### 5.2 Projection on RV space

From a mathematical point of view, equation (16) corresponds to a projection onto a lower-dimensional space, on which we can approximate the dynamics of the moments of the class-specific morph distributions by the dynamics of the moments of the RV-weighted morph distribution. For simplicity, we call this approximation the projection on RV space. Mathematical details are given in Appendix C, but the key biological insight is that, if morph distributions are sufficiently clustered around the morph mean, the dynamics of the morph moments on the slow time scale will closely track the moments of the reproductive-value-weighted morph distribution.

Equation (13) shows that the dynamics of the morph reproductive values are *O*(1), so that reproductive values change on the same time scale as class frequencies. On this fast time scale, we can derive the following quasi-equilibrium approximation from equations (10) and (13):

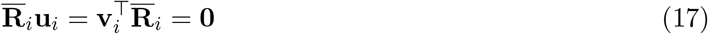

so that the vectors of class frequencies and reproductive values are respectively the right and left eigenvectors of the matrix 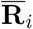 associated with eigenvalue 0, where it follows from equation (7) that 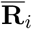 has elements 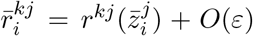. This is a multi-morph extension of a standard result from monomorphic theory (Taylor, 1990; Rousset, 2004; Lehmann & Rousset, 2014; Lion, 2018a,b; Priklopil & Lehmann, 2020).

### 5.3 Dynamics of morph means on RV space

On the slow time scale, the dynamics of the morph mean across all classes can be approximated, to leading order, by the dynamics of the reproductive-weighted morph mean (Appendix C),

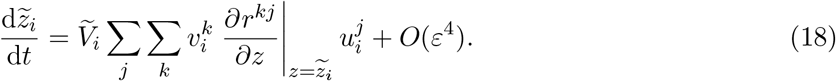

where 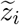 and 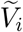 are the mean and variance of 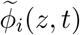. In matrix form, equation (18) can be written as:

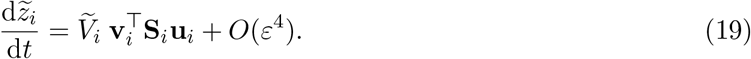

where

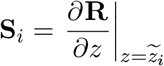

is the directional selection matrix.

The effect of selection on the morph mean is thus scaled by the morph variance, 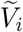, and the direction of selection is given by the selection gradient, 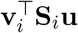 which depends on (1) the marginal effect of a change in the trait on the between-class transition rates *r^kj^*(*z*), evaluated at the morph mean, (2) the relative quality of class *k* for morph-*i* individuals, measured by the reproductive value 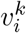, and (3) the relative quantity 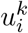 of class-*k* individuals among morph-*i* individuals. The class frequencies, morph frequencies and reproductive values are all calculated using the zeroth-order terms of the equations (8)–(11). In particular, the vectors **v**_*i*_ and **u**_*i*_ satisfy equation (17).

The selection gradient 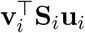 is the multi-morph extension of the classical expression for the class-structured selection gradient of invasion analyses (Taylor, 1990; Rousset, 1999, 2004; Lehmann & Rousset, 2014; Lion, 2018a,b; Priklopil & Lehmann, 2020), which can be recovered by noting that, in the single-morph case, 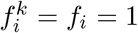, so that **u**_*i*_ = **f** and **v**_*i*_ = **v**. Equation (18) also has similarities with some quantitative genetics results (e.g. equation (8) in Barfield et al. (2011)), but with two subtle differences. First, equation (8) in Barfield et al. (2011) is derived under frequency-independence, and as a result the directional selection matrix depends on the derivatives of the mean transition rates 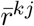 with respect to the mean trait, instead of the derivatives of the transition rates *r^kj^*(*z*) with respect to *z* (Iwasa et al., 1991; Day, 2005; Lion, 2018c). Second, equation (8) in Barfield et al. (2011) also depends on the class-specific means and is therefore unclosed (see Appendix C for further discussion).

Figure 2 illustrates the convergence to RV space in a simple two-class model. The dynamics of the morph means in classes *A* and *B* (black lines) quickly relax then closely follow the prediction of equation (18) (gray line), until the eco-evolutionary dynamics stabilise.

**Figure 2:**
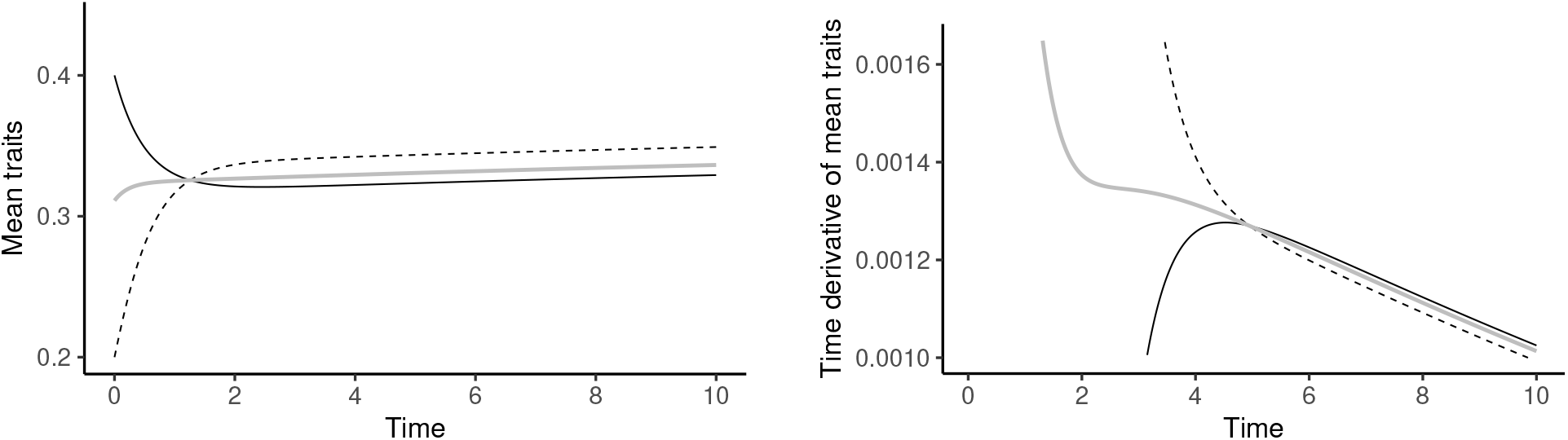
Illustration of the relaxation to RV space in a two-class model. (A) Convergence of the class-specific morph means 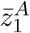 (black solid line) and 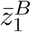 (dashed line) to the RV-weighted mean 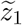 (grey line). (B) Convergence of the time derivatives 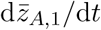 (black solid line) and 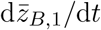 (dashed line) and shown to converge towards the prediction of the right-hand side of equation (18) (grey line). The simulation is the same as in figure S.3A, to which the reader is referred for additional details.

### 5.4 Dynamics of morph variances on RV space

As for the morph mean we can use reproductive values to derive an aggregate equation for the dynamics of the morph variances. In its most compact form, it takes the form of a Price equation (Appendix C)

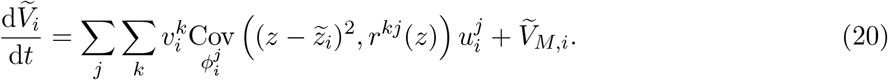

where the covariance is calculated over the distribution 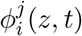. The term 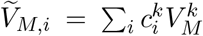 is a RV-weighted mutational variance.

The oligomorphic approximation (together with a Gaussian closure at the morph level) allows us to further expand the covariance (see equation (21) in Appendix C) as

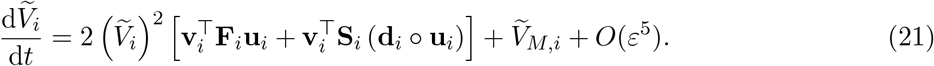

where the notation ◦ denotes the elementwise (Hadamard) product. The first-term between brackets depends on the matrix

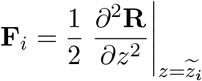

It is the class-structured analog of equation (37) in Sasaki & Dieckmann (2011) and gives the net effect of the curvature of the fitness functions *r^kj^*(*z*) on the dynamics of variance.

The second term between brackets depends on the directional selection matrix **S**_*i*_, and on the vector **d**_*i*_ of phenotypic differentiations 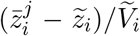. The operation **d**_*i*_ ◦ **u**_*i*_ returns a vector with elements 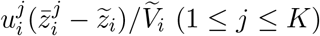. This second term represents the additional effect of directional selection on the dynamics of morph variance. Intuitively, these two effects can be understood from the decomposition of the morph variance into class-specific morph moments:

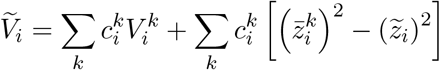

The first term shows that, when the class means are equal, the morph variance 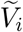 is the weighted average of the class-specific morph variances 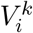. The second term shows that, even when the class variances 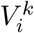 are zero, substantial differentiation between classes 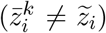 can contribute to morph variance. Hence, changes in the mean traits have a direct effect on the dynamics of the variance, which is proportional to the strength of selection in each class (captured by the matrix **S**_*i*_) and to the level of phenotypic differentiation in each class, compared to the mean morph value (captured by the vector **d**_*i*_).

Note that the second term between brackets in equation (21) still depends on the class-specific morph means, 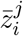. However, as shown in Lion (2018b), the vector of phenotypic differentiation is a fast variable and can be approximated, on the RV space, by a quasi-equilibrium expression which we can express solely in terms of **v**_*i*_, **u**_*i*_, 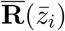 and **S**_*i*_, leading to a closed system at the morph level (Online Appendix S.2; Lion (2018b)). Although the resulting expression in the general case is not very enlightening, we show in section “Applications” and Appendix D that it yields very simple expressions when the model has only two classes.

## 6 Putting everything together

The above derivation yields a coupled system of differential equations describing how ecological densities, morph frequencies, morph means and morph variances change over time. There are three main ways to use these equations, depending on whether the biological question of interest focuses on dynamics or statics.

### 6.1 Eco-evolutionary dynamics: oligomorphic approximation

The oligomorphic approximation allows us to derive an approximation of the full eco-evolutionary dynamics that takes the form of a fast-slow system with different intrinsic time scales. We collect these equations here, for ease of reference:

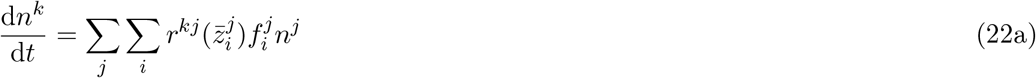

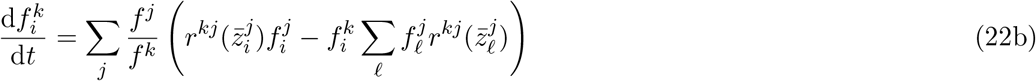

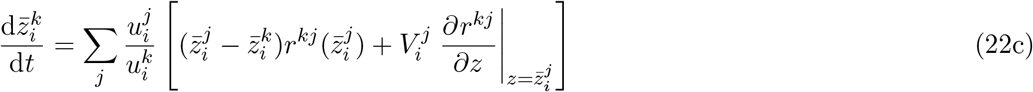

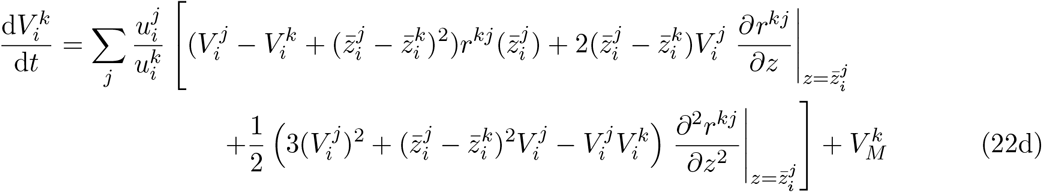

Here, we have used the Gaussian moment closure and dropped the order terms for simplificity. This system can be numerically integrated and allows us to track the joint dynamics of ecological and evolutionary variables. It also provides analytical insights into the observed dynamics. Thus, equation (22) can be viewed as an extension of classical quantitative genetics methods to take into account ecological feedbacks, class structure, multimodal distributions and the dynamics of variance. The accuracy of the approximation will tend to decrease as within-morph variation or mutation becomes too large.

### 6.2 Eco-evolutionary dynamics: projection on RV space

The projection on RV space can be used to reduce the dimension of our system and derive the dynamics of the morph means and variances:

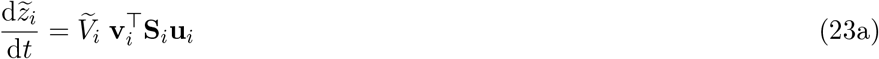

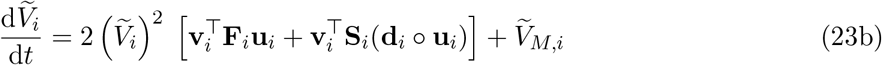

where **d**_*i*_ measures the phenotypic differentiation between classes. Both 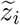 and 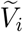 are slow variables when the morph variances are small. We can then use a quasi-equilibrium approximation, as typically done when dealing with fast-slow systems. In particular, the frequencies **u**_*i*_ and reproductive values **v**_*i*_ satisfy the following quasi-equilibrium relationships:

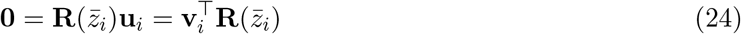

Note that, in these equations, the matrices **R**, **S**_*i*_ and **F**_*i*_ depend on the ecological densities and morph frequencies (see for instance Examples 2 and 3 below), which can be calculated from

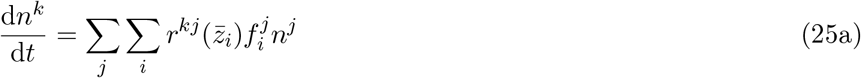

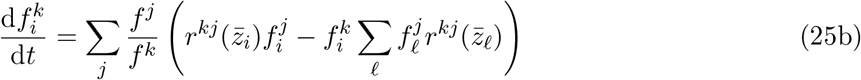

These are simply equations (22a)–(22b) where the class means 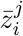 have been replaced by the morph means 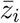. As the morph means 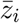 change slowly, the fast variables *n^k^* and 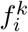 will quickly track these changes. In practice, we can either calculate, for a given value of 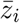, the quasi-equilibrium values of *n^k^* and 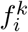 by setting the right-hand sides of equations (25) to zero, or we can numerically integrate equations (25), together with equations (23), to obtain the values of *n^k^* and 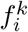 at any given time.

### 6.3 Eco-evolutionary statics

Although our framework is geared towards dynamics, it is interesting to show how we can use equations (23) to derive analytical information on the potential eco-evolutionary endpoints and their stability, and therefore recover results from invasion analyses. We then use a step-by-step analysis. First, we assume that morph means and variances change so slowly that the ecological dynamics reach an equilibrium attractor. We then plug this information into the dynamics of morph means and calculate evolutionary singularities from

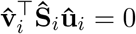

where the “hat” indicates that all these quantities are calculated on the ecological attractor. This is basically the same result as we could obtain with a traditional invasion analysis (see e.g. Taylor (1990), Rousset (2004), and Priklopil & Lehmann (2020)), but in a polymorphic resident population. As shown by Sasaki & Dieckmann (2011), we can use the dynamics of the means near these singularities to investigate their convergence stability (i.e. whether or not they attract the trait dynamics). Similarly, we can investigate the evolutionarily stability of these singularities (i.e. whether selection is disruptive or stabilising) by evaluating the dynamics of variance near these singularities. As shown by Sasaki & Dieckmann (2011), the singularities 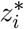 are all evolutionarily stable if and only if the morph variances converge to zero, leading to the conditions

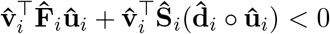

when 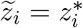. The first term was previously obtained using invasion analyses (see e.g. the first term of equation 34A in Ohtsuki et al. (2020)). The second-term captures how directional selection in the different classes contributes to disruptive selection, and probably bears some resemblance with the first term of equation 34B in Ohtsuki et al. (2020), although a precise comparison of the two results is currently beyond our reach.

## 7 Applications

As proof-of-concept examples, we apply our general framework to two-habitat migration-selection models. We start by giving general dynamical equations for these models, then explore three different biological scenarios that illustrate various aspects of our framework. From now on, we deliberately simplify the mathematical expressions by omitting the order terms and using the more intuitive notations 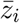 and *V_i_* to denote the (RV-weighted) morph means and variances, instead of the tilded versions.

### 7.1 General results for two-habitat migration-selection models

We consider a population inhabiting two habitats, *A* and *B*, coupled by migration. We define *m_jk_* as the migration rate from *k* to *j* and *ρ_k_*(*z*) as the growth rate of individuals in habitat *k*, which is a function of a focal trait *z*. With our notations, we have

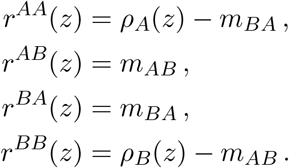

Because the migration rates do not depend on the focal trait *z*, the dynamics of morph means and variances take the following simple form using the projection on RV space (Appendix D):

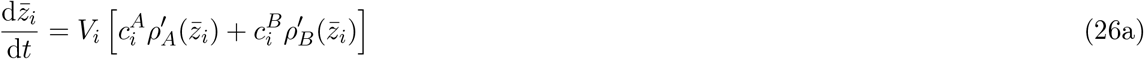

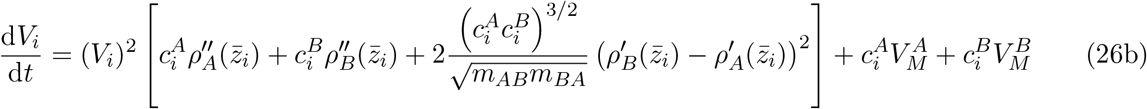

together with the following quasi-equilibrium expressions for the class reproductive values (Appendix D, Online Appendix S.4)

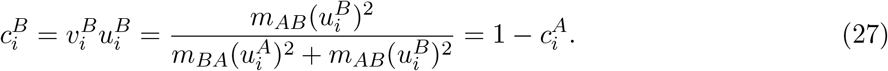

In practice, the dynamics of the densities and morph frequencies can be obtained from

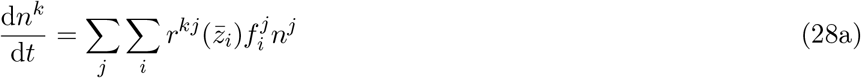

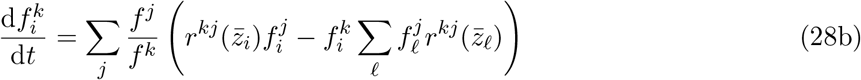

which are simply equations (25).

### 7.2 Example 1: local adaptation under mutation-selection-migration balance

In our first example, we revisit a classic model of local adaptation and investigate how migration and selection interplay to favour or hamper the generation of polymorphism in a population that can exploit two habitats with different qualities. In contrast with previous studies, we provide dynamical equations that capture the notion of habitat quality in polymorphic populations through the concept of morph-specific reproductive values we introduced in the previous sections.

We assume that, within each habitat, reproduction depends on a trait *z*. We further assume that there is a quadratic cost to fecundity and that each habitat is characterised by a value *θ_k_* which minimises the cost and that we call the habitat optimum. We thus write

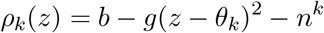

where *b* is the birth rate, *g* is the fecundity cost, *θ_k_* is the optimum in habitat *k*. The same fitness functions were used in previous studies (Meszéna et al., 1997; Ronce & Kirkpatrick, 2001; Débarre et al., 2013; Mirrahimi & Gandon, 2020) but, in contrast with most of these previous studies, we consider the possibility of asymmetric migration rates (*m_AB_* ≠ *m_BA_*).

Together with equations (23), these expressions for the vital rates allow us to derive the following equations for the dynamics of morph means and variances (Online Appendix S.6):

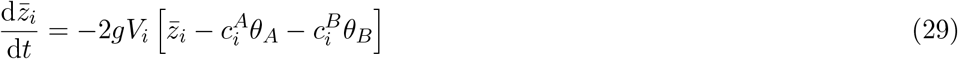

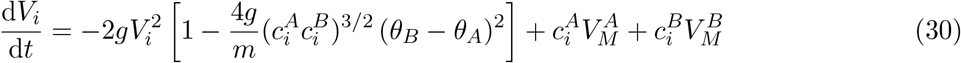

where 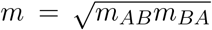 is the geometric mean of the dispersal rates, 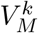 are the habitat-specific mutational variances (Appendix E), and the morph-specific class reproductive values are given by their quasi-equilibrium expressions (equation (27)). Using these equations, we investigate three main questions: (1) What are the attractors of the eco-evolutionary dynamics? (2) When does selection lead to a unimodal vs. bimodal equilibrium distribution? and (3) What is the effect of habitat-specific mutation on the evolutionary outcome? Since a full analysis of the model would be beyond the reach of this article, we focus on the most salient features in the main text and briefly consider additional technicalities in Online Appendix S.6.

#### Evolutionary attractors

From equation (29), we see that the morph means stabilise when

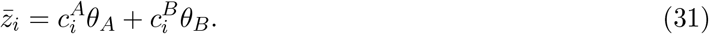

It is important to note that, because of the quasi-equilibrium approximation, the class reproductive values are also functions of 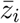, so that equation (31) is only implicit. Nonetheless, the biological implication is that the potential attractors for the dynamics of the morph means correspond to the reproductive-value-weighted mean of the habitat optima. Hence, the outcome of selection for a given morph depends on the relative difference in quality between habitats, measured by 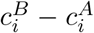. When 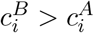, the morph mean will be biased towards the optimum of habitat *B*, whereas the opposite will be observed for 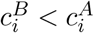. If both habitats have similar quality 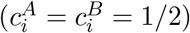, the morph mean will converge towards (*θ_A_* + *θ_B_*)/2. In the following, we fix *θ_A_* = 0 = 1 – *θ_B_*, without loss of generality.

Our analysis allows us to recover the important result that, when the mean migration rate is sufficiently high, the eco-evolutionary dynamics settle on an unimodal distribution, while bimodal distributions can be generated if the mean migration rate is below a threshold *m_c_* (figure 3A; see also Débarre et al. (2013) and Mirrahimi & Gandon (2020)). Simulations of the full model, without the oligomorphic approximation, show that both the unimodal and bimodal endpoints are accurately predicted by the projection on RV space (figure 3A).

**Figure 3:**
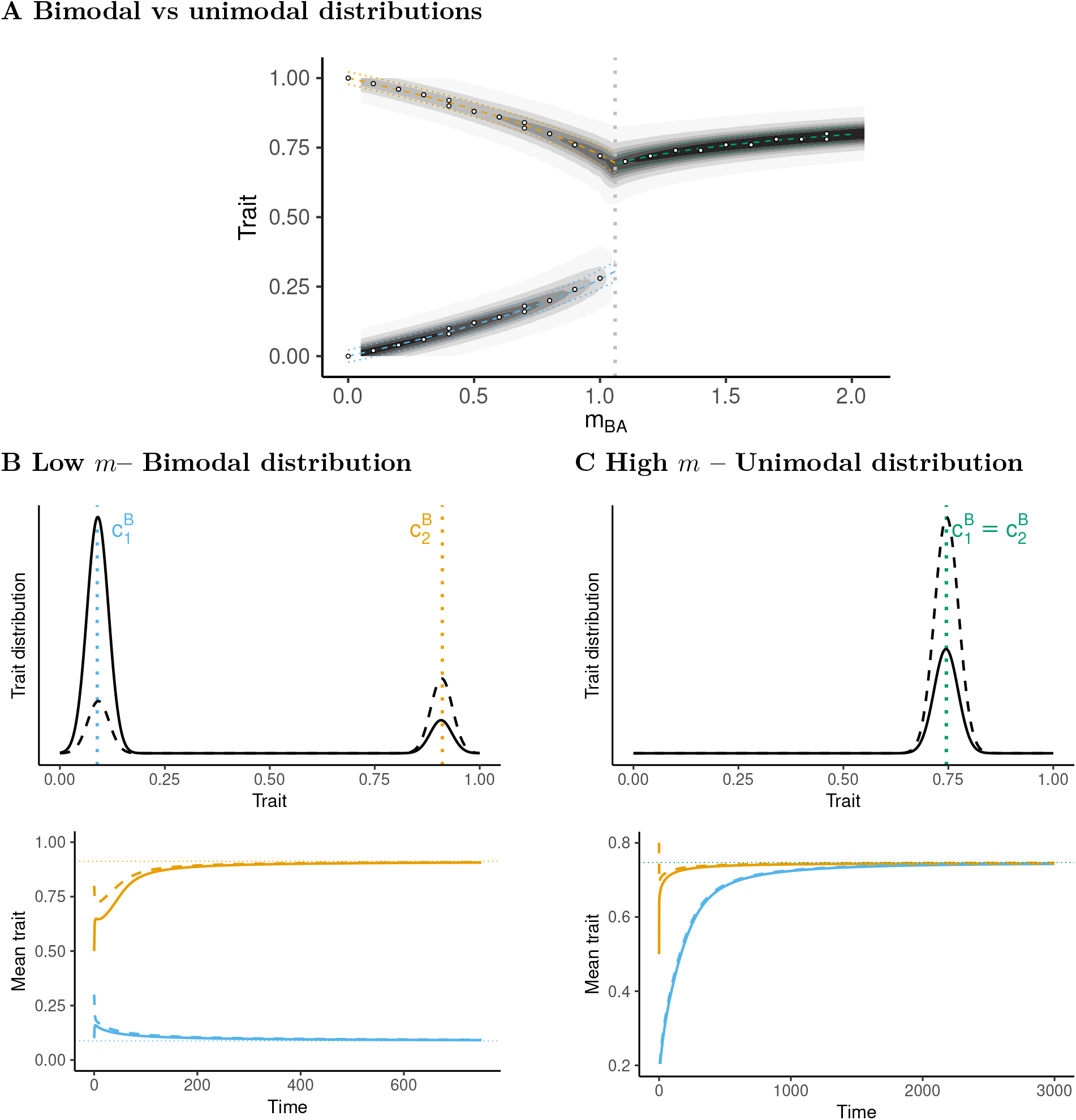
Bimodal vs. unimodal equilibrium distributions (Example 1) Figure (a) shows how the equilibrium distribution changes when *m_AB_* = 0.8 is fixed and *m_BA_* varies. For simplicity, we only show the dimorphic solution in the bistability region. The vertical dotted line represents the value *m_BA_* ≈ 1.06 at which the dimorphic equilibrium loses its demographic stability and one of the two morphs goes extinct. The analytical predictions using the projection on RV space are also shown for the mean (dashed lines) ± standard deviation (dotted lines). The white dots show the results of numerical simulations of the full model, without the oligomorphic approximation. Figures (b) and (c) show the equilibrium trait distributions and dynamics of mean traits in habitat *A* (solid line) and *B* (dashed line) obtained by numerical integration of a 2-morph oligomorphic approximation for *B_A_* = 0.4 (b) or *m_BA_* = 1.4 (c). The dotted lines indicate the value of the class reproductive value of habitat *B* for morphs 1 (blue) and 2 (orange), computed from the oligomorphic approximation. Parameter values: *b* = 1, *g* = 2, *m_AB_* = 0.8, *V_M_* = 10^-6^.

At a unimodal distribution, the class reproductive values of the different morphs are equal (e.g. 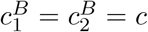 if we start with 2 morphs), so there is effectively a single morph in the population with mean trait value 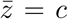 (figure 3C). In contrast, if different morphs have distinct class reproductive values (e.g. 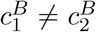), the eco-evolutionary dynamics converge towards a bimodal equilibrium distribution, where the morphs have different means (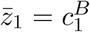 for morph 1 and 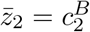 for morph 2), and occur with different frequencies in the two habitats. The trait distributions in each habitat therefore have two peaks, one around 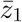 and one around 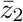, the heights of the peaks being determined by the morph frequencies 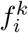 and the densities *n_A_* and *n_B_* (figure 3B). In Online Appendix S.6, we show how the positions 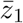 and 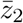 of each morph at equilibrium can be analytically calculated, thereby allowing us to recover previous results on local adapatation (Débarre et al., 2013; Mirrahimi & Gandon, 2020).

#### Disruptive selection and the balance in habitat quality

When does the interplay between migration and selection leads to unimodal vs bimodal equilibrium trait distributions? To answer this question, we turn to the dynamics of morph variances (equation (30)), which provides a useful interpretation of evolutionarily stability in terms of reproductive values. Indeed, the product 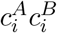 is a measure of the balance in habitat quality: it reaches its maximal value, 1/4, when the habitats have equal qualities 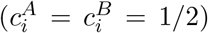 and is zero when 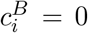 or 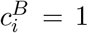. Selection on morph *i* is therefore stabilising if the balance in habitat quality is below a threshold determined by the ratio between migration (*m*) and selection (*g*):

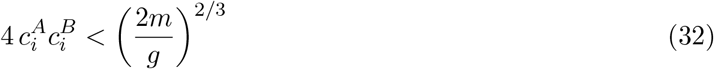

Note that, for *m* > *g*/2, this is always satisfied, but with a lower mean migration rate, stability depends on the magnitude of habitat differentiation. Hence, for a fixed *m* < *g*/2, disruptive selection is possible and may lead to the splitting of an initially unimodal distribution into two peaks, provided the habitat qualities at the monomorphic evolutonary attractor are sufficiently similar (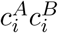 large enough). Calculating 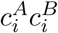 at the dimorphic attractor then shows that it is evolutionarily stable when *m* < *g*/2 and unstable otherwise. However, we note for completeness that there is a critical value *m_c_* < *g*/2 above which the bimodal equilibrium loses its demographic stability and one of the two morphs goes extinct (figure S.5A). Hence, for *m* > *m_c_*, the only possible evolutionary outcome is a unimodal distribution. For *m* < *m_c_*, multistability is possible, where the system can converge towards unimodal or bimodal distributions depending on initial conditions (Online Appendix S.6; Débarre et al. (2013)).

#### Mutation-selection-migration balance

With mutation and stabilising selection, the morph variances can settle at a mutation-selection equilibrium. From equation (30), the equilibrium variance is then given by

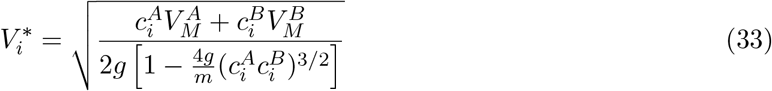

The numerator is the effect of mutation on variance, which takes the form of a reproductive-value-weighted average mutational variance, while the denominator gives the effect of selection. When only one habitat is present (e.g. 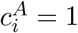 and 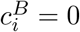), the morph variance at equilibrium is 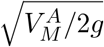, which is equation (17) in Débarre et al. (2013). For the bimodal attractor, both morphs have the same equilibrium variance if the mutational variance is the same in both habitats:

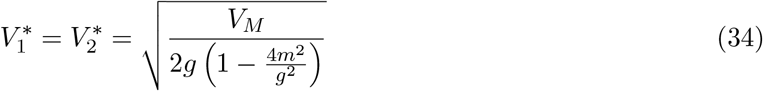

On the other hand, if 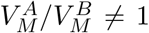, each morph has a different equilibrium variance, which can be analytically calculated (Online Appendix S.6). Figure 4A shows that this accurately predicts the equilibrium values of morph variances under mutation-selection balance.

**Figure 4:**
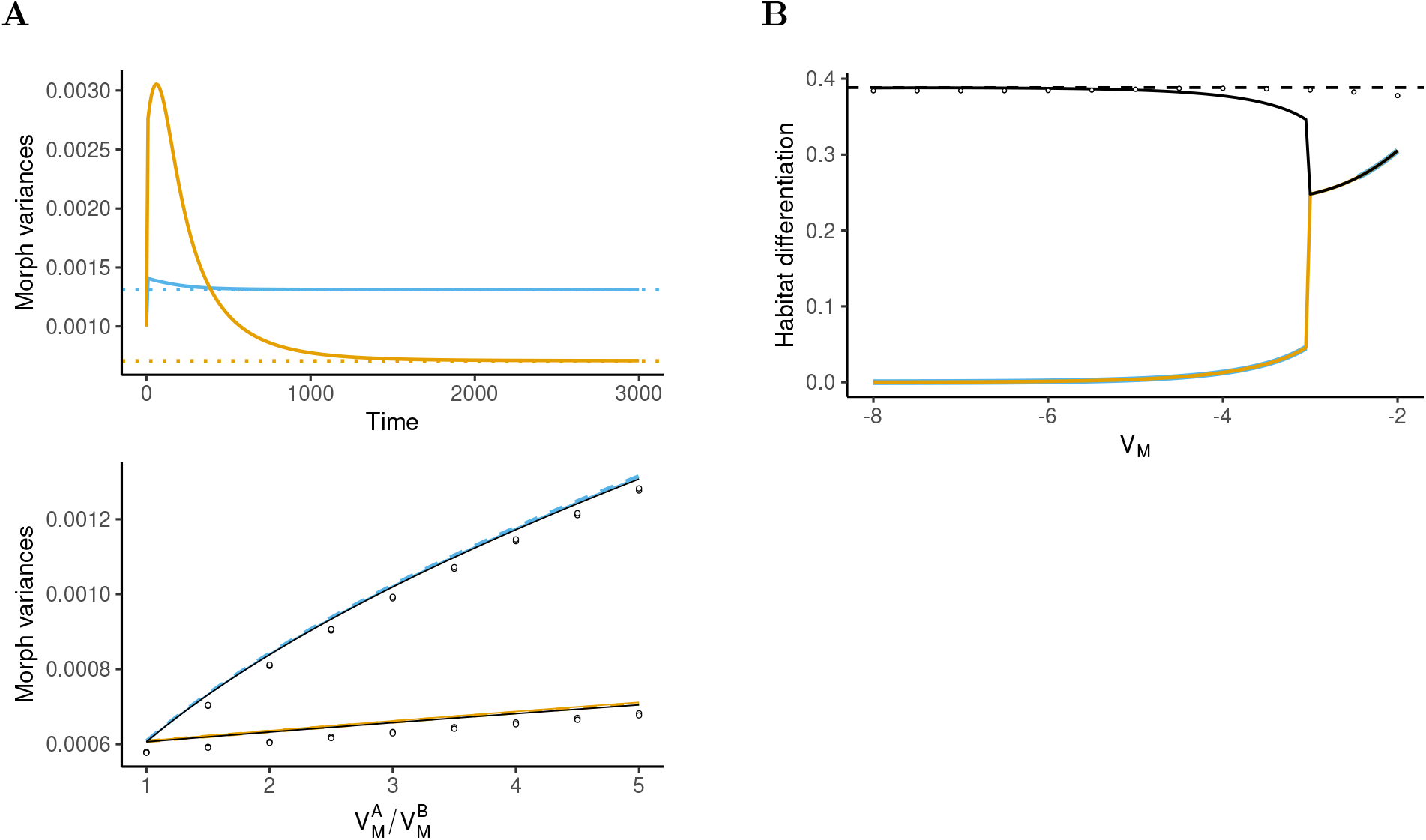
Effect of mutation (Example 1) (a) Effect of habitat-specific mutation. The top panel shows the dynamics of the variances of morph 1 (blue) and 2 (orange) which converge to mutationselection equilibrium values, which are accurately predicted by the reproductive-value-weighted formula (33) (dotted lines) with 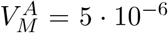 and 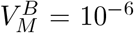. The bottom panel shows the effect of the ratio 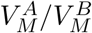 on the variances 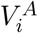 (solid line) and 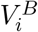 (dashed line) for morphs 1 (blue) and 2 (orange). The white dots give the results of simulations of the full model without the oligomorphic approximation. (b) Effect of the magnitude of mutational variance on the mean habitat differentiation 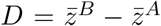 at equilibrium (black line). Here, we use 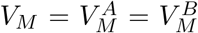. The dashed line gives the analytical prediction of equation (S.41), the white dots the simulations results, the blue and orange lines give the morph-specific habitat differentiation 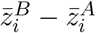. Parameters in all panels: *b* =1, *g* = 2, *m_AB_* = 0.8, *m_BA_* = 0.4.

As mutation increases, the accurary of the oligomorphic approximation is expected to decrease, as high mutation will tend to generate broader morph distributions. Nonetheless, figure 4B shows that our analytical prediction of the differentiation 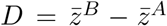 remains good even for relatively large values of the mutation variance (see Online Appendix S.6 for a more detailed discussion). The approximation breaks down roughly when the mutation variance is of the same order as the morph variances.

### 7.3 Example 2: evolution of intra-specific competition

In our second example, we assume that, within each habitat, competition between individuals depends on the competitive ability *z*. As previously, we further assume that there is a quadratic cost to competitiveness and that each habitat is characterised by a value *θ_k_* which minimises the cost. We thus write

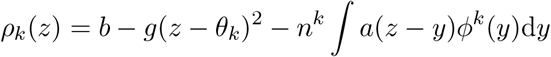

where the competition kernel *a*(*z* – *y*) represents the effect of competition by an individual with trait *y* on an individual with trait *z*. Importantly, the fitness functions *ρ_k_* now depend on the distributions *ϕ^k^*(*y*, *t*), in contrast to our Example 1. Following Sasaki & Dieckmann (2011), we decompose this distribution into a sum of the distributions *ϕ_ℓ_*(*y*, *t*) and, for each of these distributions, we use a Taylor expansion of the competition kernel near 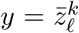 to express the competition experienced by a focal individual in terms of the means of all the other morphs. We obtain (Online Appendix S.7)

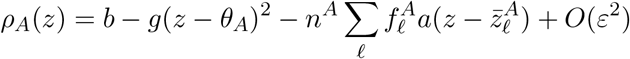

and a similar expression for the growth rate in habitat *B*. We can use these expressions to calculate the quantities 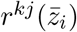, and the partial derivatives evaluated at 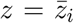, and plug these expressions in equation (22) to obtain the final approximation for this model (Online Appendix S.7).

This example is a perfect illustration of how all the equations of system (22) are coupled. In particular we have:

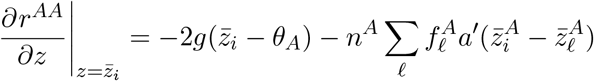

which shows that the competition experienced by morph *i* depends on the other morphs through their frequencies 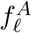 and mean trait values 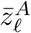. Hence, the dynamics of the mean of morph *i* depends on the frequencies and mean traits of the other morphs.

A full analysis of the model is beyond the scope of this paper, but we show in figure 5 the dynamics for a specific choice of parameters. Figure 5C shows that, starting from an effectively monomorphic population (where the two morphs have approximately the same mean values), disruptive selection leads to the splitting of the population into a bimodal distribution after *t* ≈ 1000. Disruptive selection is indicated by the explosion of morph variances at the same time (figure 5D). The population then stabilises around a dimorphic equlibrium distribution with means 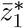 and 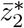. Stabilising selection is indicated by the decrease in variance after branching. Figure 5B further shows that the morphs have different frequencies in each habitat (i.e. morph 1 is slightly more abundant in habitat *B* while morph 2 is slightly more abundant in habitat *A*), although they have similar values for the means and variances in both habitats.

**Figure 5:**
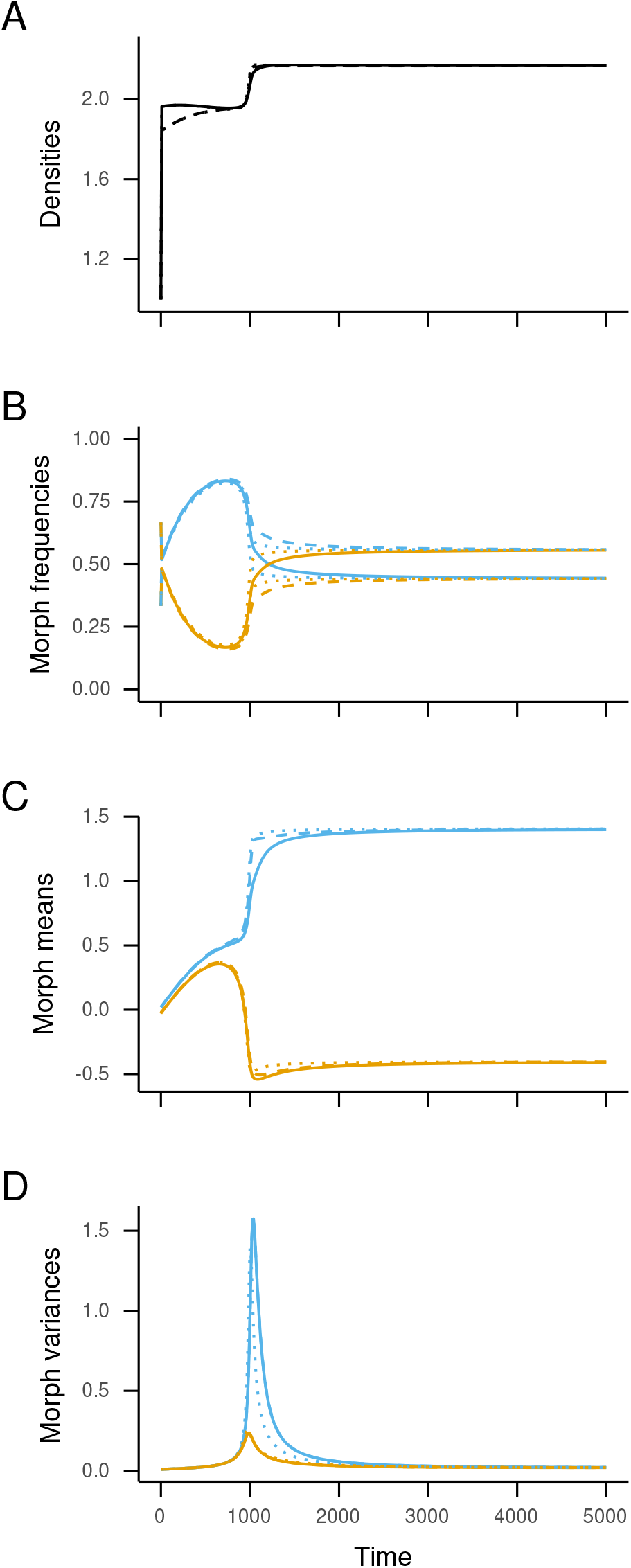
Eco-evolutionary dynamics of the resource competition model (Example 2) (A) Dynamics of densities of individuals *n^k^* in habitat *A* (solid line) and *B* (dashed line). (B) Dynamics of the morph frequencies 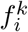 in habitat *A* (solid line) and *B* (dashed lines) for morph 1 (blue) and 2 (orange). (C) Dynamics of the morph means 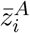 (solid lines) and 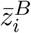 (dashed lines). (D) Dynamics of the morph variances 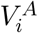 (solid lines) and 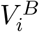 (dashed lines). In all panels, the results of a numerical integration of equations (22) are shown and the corresponding predictions of the RV projection (26)–(28) are overlaid as dotted lines. Parameters: *b* = 1, *g* = 0.1, *θ_A_* = 0 = 1 – *θ_B_*, *m_AB_* = *m_BA_* = 0.3, *a*(*x*) = 0.5 exp(−*x*^2^/8), *V_M_* = 10^-5^.

This latter observation suggests that it may be interesting to find a simplified description of the population at the morph level, by aggregating habitat-specific morph moments into a single measure. This is precisely the goal of the RV projection, and we show in Online Appendix S.7 that we obtain the following expression for the dynamics of morph means:

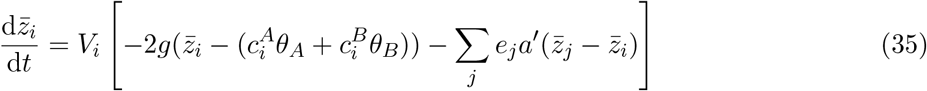

where *e_j_* measures the average net fitness effect on individuals of morph *j* due to competition with the other morph. For morph 1, we have 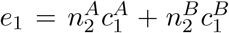, and the expression for *e*_2_ is obtained by swapping 1 and 2 subscripts. This expression shows that the class reproductive values give the proper weights to measure the intensity of competition in each habitat. Figure 5 shows that the RV projection (dotted lines) accurately predicts both the trajectories end endpoints of the densities, morph frequencies, morph means and morph variances.

Equation (35) tells us that, starting from a quasi-monomorphic situation (i.e. 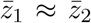, so that 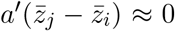), the population will converge towards the RV-weighted mean of the habitat optima, 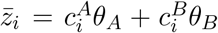. However, at this point selection becomes disruptive and a bimodal equilibrium distribution is eventually reached, with peaks located at the equilibrium morph means. From equation (35), the morph means satisfy:

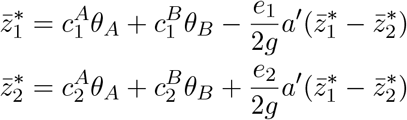

This provides an intuitive interpretation of the position of the two peaks of the equilibrium distribution as the deviation from the morph-specific mean habitat optimum due to competition with the other morph. In the single-class case, we recover the results of Kisdi (1999) (see e.g. her equation (10)).

### 7.4 Example 3: transient out-of-equilibrium evolution in a resource-consumer model

In our final example, we show how our approach can be used to analyse eco-evolutionary dynamics across different time scales. We consider a resource-consumer model where the resource is produced at different rates in the two habitats, so that *S^k^*(*t*) is the density of resource in habitat *k* at time *t*. We focus on the evolution of a consumer’s exploitation rate, *z*, and assume a type-II functional response such that

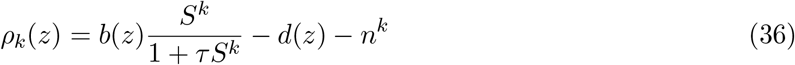

where *b*(*z*) is the fecundity rate, *d*(*z*) is the mortality rate, and *τ* depends on the handling time. The dynamics of the resources *S^A^* and *S^B^* are given in Online Appendix S.8. In our thought experiment, we introduce the same low density of consumers in each habitat, but the genetic composition of each subpopulation is different: the initial distribution in each habitat is bimodal, with two peaks representing more prudent vs more rapacious consumers, but habitat *A* is predominantly seeded with more prudent consumers, while habitat *B* is predominantly seeded with more rapacious consumers (figure 6A). Hence, the mean value of the trait in habitat *A* is initially lower than in habitat *B*.

**Figure 6:**
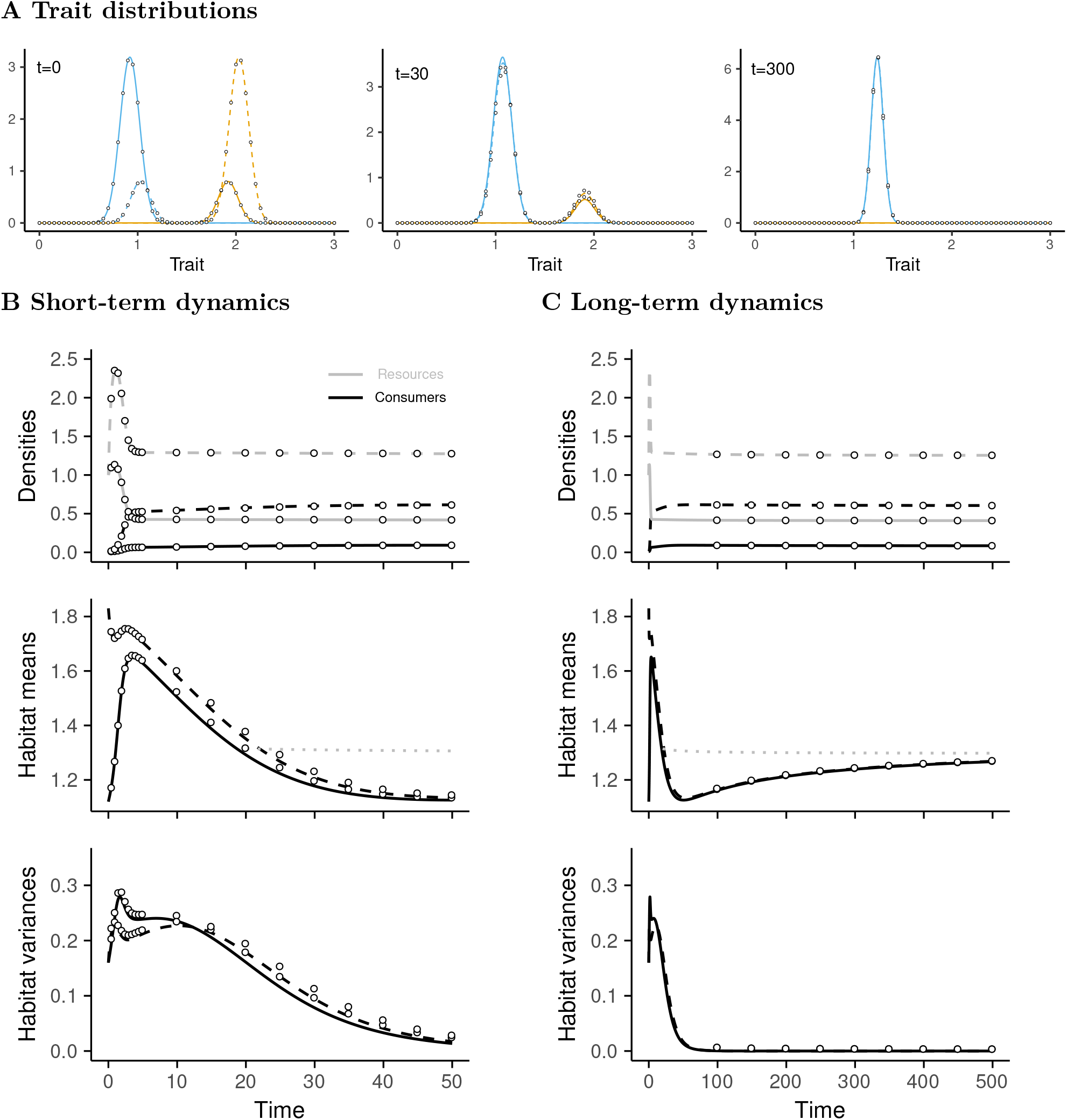
Eco-evolutionary dynamics of the resource-consumer model (Example 3) Panel (A) gives the distributions in each habitat at different times (A=plain line, B=dashed line, morph 1= blue, morph 2 = orange). Panel (B) and (C) give the dynamics of densities *n^A^* and *n^B^* (the consumer, black) and resources *S^A^* and *S^B^* (grey), habitat means 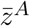 and 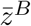 and habitat variances (plain line: habitat *A*; dashed line: habitat *B*). Panels (B) and (C) show the same simulation results, but on different times scales (*t = 50* in (B), and *t = 500* in (C)). In all panels, the main results are obtained using the RV projection, and the dots show the results of the numerical simulations of the full model without the oligomorphic approximation. The horizontal dotted grey line is the prediction of the ESS 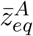 (equation (39)). Parameter values are given in Online Appendix S.8.

Our aim is to track how the habitat-specific means 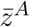 and 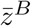 change over time, much like a field quantitative geneticist would measure time series for phenotypes at different locations. In terms of the morph frequencies and morph means, we have 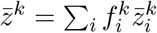 and, together with our projection on RV space, this allows us to derive the following equation (Online Appendix S.8)

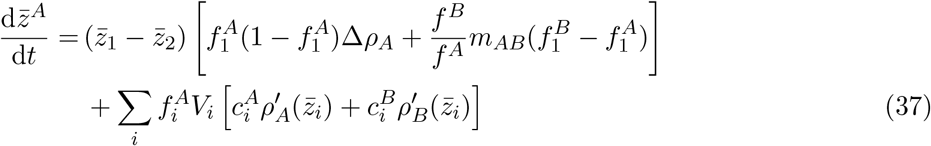

where 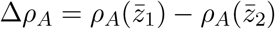 is the average difference in growth rates between the two morphs.

Equation (37) is particularly enlightening about how the different time scales of eco-evolutionary dynamics interplay. The first line of equation (37) represents the contribution of the dynamics of frequencies to the change in the habitat-specific mean (i.e. what happens when the heights of the peaks change), while the second line represents the contribution of the change in morph means (i.e. when the position of the peaks change). When the distance between the two morphs 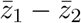 is large compared to the within-morph variances *V_i_*, the dynamics of the mean trait in habitat *A* is dominated by the change in frequencies. The first term between brackets is reminiscent of classical population genetics equations and tells us that the dynamics of the mean trait in habitat *A* depends on the difference in growth rates between the two morphs, scaled by the variance 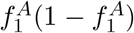. The second term between brackets captures the effect of migration from *B* to *A* and reveals that 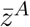 will tend to increase due to migration if the morph with the larger trait value is more abundant in habitat *B*. Thus, the first line in equation (37) captures the interplay between selection and migration typical of classical population genetics (or two-species ecological) models, but in contrast with these approaches it does not assume that there is no standing variation within a morph.

On the other hand, when the morph means are close (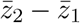 is small, so that the population can be thought of as quasi-monomorphic), the dynamics of the habitat mean is dominated by the change in morph means, which is described by the second line of equation (37). The term between brackets is now reminiscent of classical expressions for the selection gradient in class-structured populations, and the contribution of each morph now depends on the slopes of the morph’s growth rates in each habitat.

Hence, if morph 1 has a higher growth rate but a lower slope than morph 2, it can be transiently selected in our model. In figure 6, we show that, starting with an asymmetric bimodal distribution at time *t* = 0 (figure 6A), the model exhibits an increase in resource abundance, due to resource production, while the consumer densities are still low. Note that the model assumes that resource production is higher in habitat *B*. During that initial phase, the mean trait increases in habitat *A* and decreases in habitat *B*. In this phase, the dynamics of 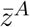 is dominated by the first line in equation (37). Once the peak in resource abundance is reached, resource consumption brings the densities of consumers up, and the resource abundance down to habitat-specific equilibrium values. As the resource abundance decreases, the selective advantage of the more prudent morph increases, and the rapacious morph is eventually competitively excluded in both habitats, as shown by the decrease in habitat variances. At this point, the term between brackets in the first line of equation (37) becomes zero, and the dynamics of 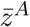 are then entirely driven by the second line. The population is now quasi-monomorphic and we have

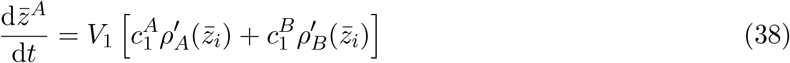

so that the mean trait increases in the direction of the slopes of the growth rates, weighted by the class reproductive values in each habitat. In our simulations, we assume *d*(*z*) = 1 + *z* and *b*(*z*) = *b*_0_*z*/(1 + *z*), which leads to the following relationship for the equilibrium value:

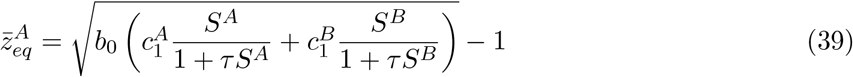

This long-term equilibrium value is indicated by the gray dotted lines in figures 6B and 6C, and we see that the slow increase of the mean trait towards this value unfolds on a much longer time scale than the fast ouf-of-equilibrum dynamics. Note that, in figure 6, the results of simulations of the full model are also given (dots), showing that our approximation very accurately captures the short- and long-term dynamics of the ecological variables and of the habitat-specific moments.

Figure 6 assumes that the initial standing variation in the population is large. This corresponds to what is called strong selection in invasion analyses (i.e. the two morphs have different means). It is interesting to compare the results with the dynamics predicted when the standing variation in the population is small, that is when the morph means are initially close (e.g. weak selection). In this case, figure S.8 shows that, following a negligible transient increase, the mean trait in each habitat slowly decreases until the equilibrium value is reached. This is easily understood from equation (37), because when 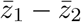 is initially small, only the slow dynamics captured by the second line drive the dynamics of the habitat mean.

Although this rapid analysis of a toy model is by no means an extensive exploration of its behaviour, it illustrates the value of our approach when analysing eco-evolutionary dynamics across different time scales. In particular, equation (37) bridges the gap between fast evolutionary dynamics in nonequilibrium population (typically analysed in ecological and quantitative genetics models) and slow evolutionary dynamics in populations that have reached an ecological attractor (typically analysed using adaptive dynamics).

## 8 Discussion

We have developed a novel theoretical framework to model the eco-evolutionary dynamics of polymorphic, class-structured populations. Our approach leads to dynamical equations that allow us to bridge the gap between quantitative genetics approaches, which are typically interested in short-term transient dynamics under substantial standing variation, and adaptive dynamics approaches, which focus on long-term eco-evolutionary statics under mutation-limited evolution. Our analysis makes two key contributions. First, we extend the recently developed oligomorphic approximation (Sasaki & Dieckmann, 2011) to class-structured populations. Since class structure is a major feature of natural biological populations, this allows the method to be applied to a broad range of ecological scenarios, taking into account individual differences in state including age, spatial location, infection or physiological status, and species. Second, we combine this approximation with recent theory on reproductive values to obtain a lower-dimensional approximation of the eco-evolutionary dynamics of multi-morph structured populations. The combination of these two approximations allows us to obtain compact analytical expressions for the dynamics of multimodal trait distributions in structured populations under density- and frequency-dependent selection. These analytical results are biologically insightful as they highlight how the quality and quantity of individuals in different classes affect the eco-evolutionary dynamics.

At a general level, our theoretical framework lies at the intersection of population genetics, quantitative genetics and adaptive dynamics. First, as in population genetics, it predicts how the frequencies of interacting morphs change over time, but explictly takes into account eco-evolutionary feedbacks. Second, we also describe the dynamics of the mean and variance of the trait distribution of each morph. This is reminiscent of moment methods typically used in quantitative genetics (Barton & Turelli, 1987; Turelli & Barton, 1990; Barton & Turelli, 1991), but our approach effectively extends these tools to multimodal distributions, frequency-dependent selection, and a broad range of ecological scenarios. Third, while our result for the dynamics of the mean takes the form of Lande’s univariate theorem (Lande, 1976, 1979, 1982; Barfield et al., 2011), we also track the dynamics of genetic variance and describe how initially unimodal distributions can split into different modes due to frequency-dependent selection. This effectively bridges the gap between quantitative genetics and adaptative dynamics to provide a more complete understanding of the evolution of quantitative traits.

Our approach broadens the scope of classic quantitative genetics theory to examine the dynamics of multimode distributions and a wider range of ecological feedbacks. Classical quantitative genetics methods are typically restricted to unimodal character distributions. However multimodal distributions are a very common outcome when frequency dependence causes disruptive selection, as often occurs when ecological feedbacks are taken into account in evolutionary models. The possibility to handle disruptive selection has been a landmark of adaptive dynamics, but it relies on the assumption that evolution is mutation-limited, so that the only source of variation in the population comes from rare mutations. Our approach relaxes these assumptions and allows us to describe the joint effect of mutation and substantial standing variation on the eco-evolutionary dynamics. Although adaptive dynamics provides tools to study evolution in polymorphic resident populations (e.g. after branching, Geritz et al. (1998), Kisdi (1999), and Durinx et al. (2008)), the resulting analysis is often difficult and restricted to potential evolutionary endpoints. In contrast, our dynamical approach makes it possible to track the joint dynamics of ecological densities and of multimodal trait distributions.

While these technical advances were already present in Sasaki & Dieckmann (2011)’s original paper, our extension to class structure makes our analysis directly applicable to a broad range of biological scenarios where individuals differ because of non-genetic factors such as age, physiological status, or spatial location. A drawback of this increased realism is that it inflates the number of ecological and genetic variables we need to track. We therefore apply recent theory on reproductive values (Lion, 2018a,b; Priklopil & Lehmann, 2020) to simplify the oligomorphic analysis and obtain a compact description of how the morph-level trait distributions change over time when there are demographic transitions between classes. The key idea is to define a weighted trait distribution that gives us a way of examining how selection acts on a particular morph across all the classes (Fisher, 1930; Taylor & Frank, 1996; Frank, 1998; Rousset, 2004; Lehmann & Rousset, 2014; Lion, 2018a). For the dynamics of the mean trait of a morph, the result takes the form of the structured extension of Lande’s theorem (see also Barfield et al. (2011)), but we also provide a new description of the dynamics of morph variances. Application of the method to the simplified two-class case gives insight into how stabilizing and disruptive selection are impacted by class frequencies and reproductive values and shows when directional selection will impact disruptive selection. As such we provide a very general tractable framework for a more complete eco-evolutionary analysis of class-structured models.

Reproductive value is an important concept in evolutionary ecology, which is ubiquituous whenever one has to deal with structured populations and provides a biologically intuitive measure of quality (Fisher, 1930; Rousset, 1999, 2004; Grafen, 2006; Lehmann & Rousset, 2014; Gardner, 2015; Lion, 2018a). As such, a limitation of previous analyses of two-habitat migation-selection models was that they did not make a clear connection between the predictions and the concept of reproductive value. Our Example 1 thus provides a conceptually useful complement to the analyses of Meszéna et al. (1997), Ronce & Kirkpatrick (2001), Débarre et al. (2013), and Mirrahimi & Gandon (2020) as it provides clear interpretations of directional and disruptive selection in terms of the reproductive values of the two habitats. We also show that our reproductive-value weighted approximation accurately predicts the numerical simulations of the full system, with or without mutation-selection balance. In addition, we show that the condition for disruptive selection takes a very simple form and can be summed up through a measure of habitat differentiation that depends on the class reproductive values. Nonetheless, it would be particularly interesting to couple our approach with the Hamilton-Jacobi framework introduced by Mirrahimi & Gandon (2020), which accurately predicts the equilibrium trait distribution even when the mutation rate is large but does not track the dynamics of the distribution over time.

An other important implication of our results is that they allow us to analyse the eco-evolutionary dynamics across different time scales. As such, our approach is very relevant to the current revival of interest on the time scales of ecological and evolutionary processes, as it can be used to examine the role of ‘fast evolution’ when this is fueled by a large standing variation at the population level. In our equations, this corresponds to morphs with very different trait means. As shown in our Example 3, our analysis allows us to shed light on the interplay between fast and slow eco-evolutionary processes, and to examine the evolutionary consequences of non-equilibrium dynamics. This has broad implications, as rapid or short-term evolution is a crucial process in conservation biology or epidemiology. For instance, our resource-consumer example predicts transient selection for increased resource exploitation, which is similar to a classic prediction that virulent pathogen strains should be favoured at the start of an epidemic and more prudent strains be selected for at endemic equilibrium (Lenski & May, 1994; Day & Proulx, 2004; Day & Gandon, 2007; Berngruber et al., 2013). Hence, our approach could be used to generate predictions for the evolutionary consequences of non-equilibrium processes such as disease seasonality (Altizer et al., 2006; Ferris & Best, 2018; Lion & Gandon, 2021), or repeated epidemics driven by antigenic escape (Sasaki et al., 2021). More generally, extensions of our approach could be used to better understand how environmental fluctuations affect disruptive selection and polymorphism, as this represents a fundamental but currently understudied research area (but see Svardal et al. (2015) for an adaptive dynamics treatment).

Our formalism has strong links with the current interest in clarifying the connection between adaptive dynamics and quantitative genetics models (Abrams, 2001; Day, 2005; Lion, 2018c) and analysing eco-evolutionary dynamics at different time scales (see e.g. Bassar et al. (2021)). This has led to various theoretical developments that employ very similar ideas to those we use here, and focus on fundamental ecological questions such as transient dynamics (Day & Proulx, 2004), community stability (Barabás & D’Andrea, 2016), multivariate traits (Mullon & Lehmann, 2019), demographic stochasticity (Débarre & Otto, 2016) or periodic environments (Lion & Gandon, 2021). Compared to these other works, a key advantage of the morph decomposition we employ is that it makes it easier to consider non-Gaussian distributions at the population level, by approximating the population-level moments in terms of morph moments, but it would be particularly interesting to couple our approach with these other technical frameworks.

At a biological level, we expect our approach will allow us to deepen our understanding of the processes that generate and maintain diversity in traits. Although the number of morphs can be arbitrarily chosen, it makes sense to use biological principles to guide this choice. For instance, in our two-habitat model, we typicaly expect that at most 2 different morphs will coexist due to the competitive exclusion principle. In practice, a scenario where the number of morphs is unsufficient can be quickly identified in numerical simulations as it corresponds to a situation where at least one morph variance diverges to infinity. At a more conceptual level, the number of morphs in our approach is linked to the dimension of the environmental feedback, which sets an upper limit to diversity (Meszéna et al., 2006; Metz et al., 2008; Lion & Metz, 2018).

We illustrate the application of the approach using two-habitat migration-selection models, but this can be broadened to examine fundamental processes such as more general forms of stage structure, species interactions, and speciation. For instance, the approach can be used to determine the time required until a population diversifies under frequency-dependent disruptive selection, which, for asexually reproducing species, is the waiting time until adaptive speciation. Additional insights will come from both ecological extensions – such as assortative mating – and in particular genetic extension – such as multi-locus inheritance, recombination, diploidy, and random genetic drift. In addition, while we have focused on the evolution of single traits, an important extension of our work would consider multi-variate traits (see e.g. Mullon & Lehmann (2019) for a quantitative genetics treatment without class structure, and Sasaki et al. (2021) for a two-trait extension of the oligomorphic approximation). In particular, plastic traits (i.e. traits that are not expressed in all classes) could be relatively easily taken into account in a multi-variate extension of the framework we present here.

To sum up, we think our analytical approach will allow for a better understanding of the role of ecological feedbacks, frequency- and density-dependent selection in nature, and has the potential to facilitate a tighter integration between eco-evolutionary theory and empirical data. At a technical level, our approach moves the field on from either focusing on unimodal character distributions, often taken in models of quantitative genetics theory, or on negligible within-morph variance, which is often assumed in models of adaptive dynamics. At a biological level, it has considerable potential to advance our understanding of the ecological factors driving the evolution and maintenance of diversity, which remains an important empirical and theoretical challenge in multiple fields and contexts.

## Acknowledgements

We thank two anonymous reviewers for very constructive and detailed comments. This study was supported by ANR JCJC grant ANR-16-CE35-0012-01 to SL, grants R01 GM122061-03 and NSF-DEB-2011109 to MB, and the ESB Cooperation Program, The Graduate University for Advanced Studies, SOKENDAI.

## Appendix A: Oligomorphic approximation

From equation (9) in the main text, it is straightforward to derive the dynamics of the within-class trait distributions *ϕ^k^*(*z*, *t*) = *n^k^*(*z*, *t*)/*n^k^*(*t*). We obtain, dropping the dependency on time for simplicity

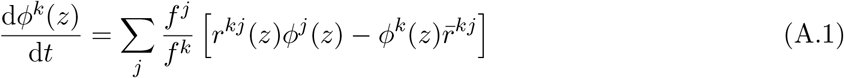

To derive the oligomorphic approximatin for our class-structured population model, we proceed as in equation (6) of Sasaki & Dieckmann, 2011 and define the dynamics of the frequencies 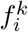 so that individuals with phenotype *z* have the same per-capita growth rate in a given class, independently of which morph they belong to. This translates into

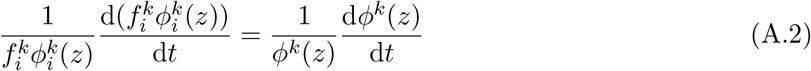

which can be rewritten as

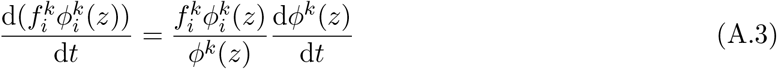

Integrating over *z* and using equation (A.1) yields

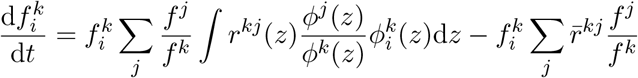

To simplify this equation, we use the following equality, which simply means that if individuals with phenotype *z* are, say, twice more abundant in class *j* than class *k*, this ratio is preserved irrespective of the morph to which they belong:

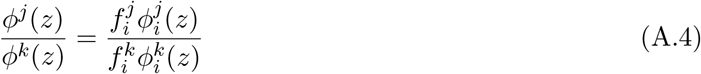

In Online Appendix S.1, we show that (A.4) naturally follows from assumption (A.2). Plugging equation (A.4) ino the dynamics of frequencies above then leads to the first part of equation (11):

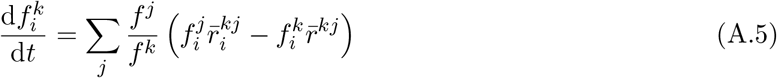

With only one class, we recover equation (7) in Sasaki & Dieckmann (2011). The second part of equation (11) in the main text can then be obtained by plugging the approximations (6) and (7) into equation (A.5).

Similarly, to calculate the dynamics of 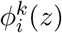, we rearrange equation (A.2) as

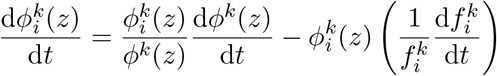

Using equations (A.1), (A.4) and (A.5) then yields after some rearrangements:

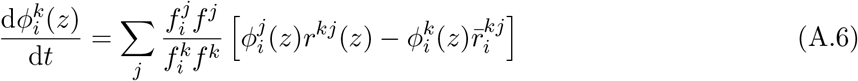

With a single class, we recover equation (8) in Sasaki & Dieckmann (2011).

## Appendix B: Dynamics of class-specific morph moments

Equation (A.6) can be multiplied by *z* or 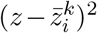 and integrated to obtain the dynamics of the morph means and variances respectively. For the morph means, we obtain:

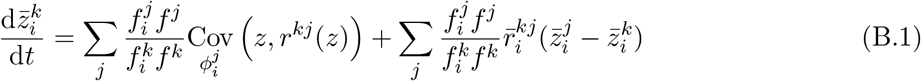

where the covariances are taken over the distributions 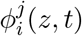 and take the form

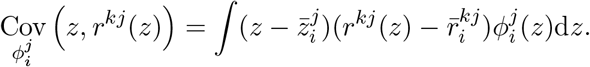

Equation (B.1) is a morph-specific version of equation (3) in Lion (2018a). Using the Taylor expansion (6) in the main text, we have

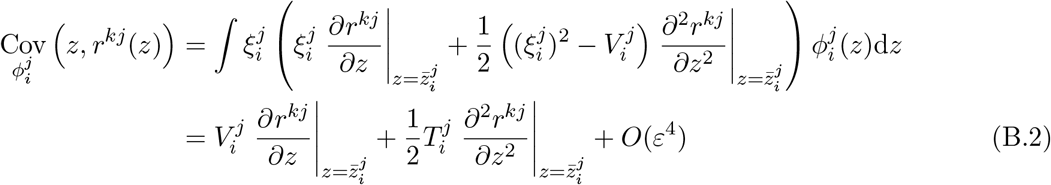

where 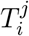 is the third central moment of 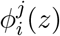, which we neglect in the following (assuming 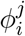 is symmetric). Substituting equations (B.2) and (7) into equation (B.1), we then obtain:

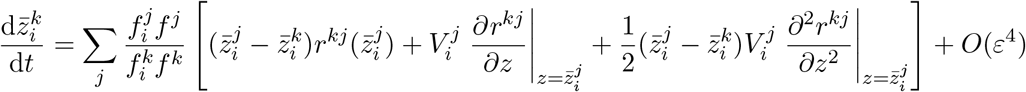

Keeping only terms up to second order in *ε*, we obtain equation (14) in the main text.

For the morph variances, defined as 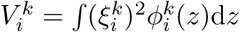, we have, using equations (A.6), (6) and (7),

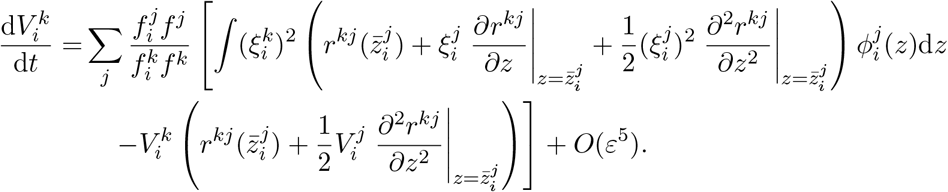

We can further simplify with the following relationships:

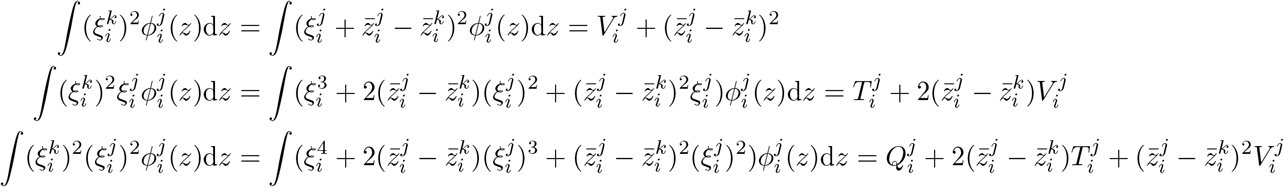

This yields equation (15) in the main text, again using the assumption that the morph distribution 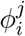 is symmetric so that 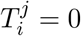.

## Appendix C: Dynamics of population-level morph moments

The derivations in appendix B yield dynamical equations for the class-specific morph moments. Depending on the question of interest it may be useful to focus on the population-level morph moments, averaged over all classes. This can be done in two ways.

### Unweighted morph distribution

A natural way is to introduce the following average distribution, which corresponds to the unweighted arithmetic mean of the class-specific distributions (Lion, 2018a):

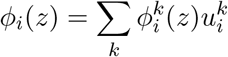

where 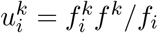 and 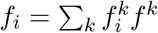 is the total frequency of morph *i* in the population. Integrating over *z* leads to a relationship between morph means: 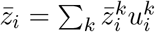, from which the following equation can be derived

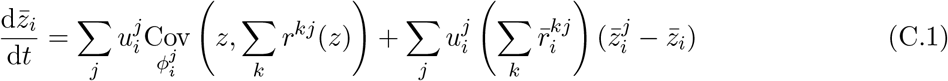

This is the morph-specific equivalent of equation (2) in Lion (2018a), and an oligomorphic approximation of equation (C.1) can be derived using approximation (6) and (7). A similar equation can be derived for the dynamics of the morph variance, 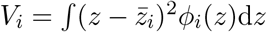.

### RV-weighted morph distribution

In equation (C.1), the dynamics of 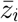 depends on the moments of the class-specific distributions 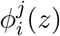. Our goal here is to find a meaningul way to summarise the dynamics using only the moments of the population-level morph distribution *ϕ_i_*(*z*).

There are two equivalent ways to do this. The first approach applies the method of Lion (2018b) at the morph level, and uses a quasi-equilibrium approximation of the phenotypic differentiations 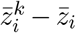 to simplify equation (C.1). This is summarised in Online Appendix S.2. The second, simpler approach is to calculate the moments of the reproductive-value-weighted distribution, as in Lion (2018a), but applied at the morph level. The time-dependent reproductive values satisfy equation (13), and the RV-weighted morph distribution is 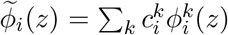 where 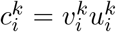. Following Lion (2018a), we obtain the following Price equation:

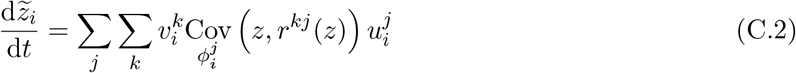

Equation (C.2) is valid irrespective of the shape of the morph distribution. Comparing equations (C.1) and (C.2) shows that, when there is no covariance between the trait and the transition rates (i.e. when the covariance terms are zero), the RV-weighted avearge 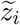 does not change while the unweighted average 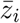 can still change due to between-class transitions. Hence, RV-weighting allows us to capture the notion that there should be no evolutionary change in a neutral model. Moreover, the dynamics of 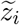 are always *O*(*ε*^2^), so that 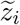 is a slow variable.

If the morph distribution is sufficiently narrow, we can approximate the covariance terms in equations (C.1) and (C.2) using equation (B.2). Both the QE and RV approaches then lead to the following equation

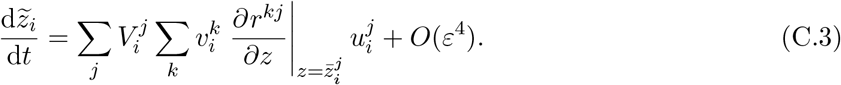

Hence, for narrow morph distributions, the morph mean and the RV-weighted morph mean have the same dynamics on the slow manifold characterised by 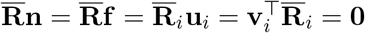.

Equation (C.3) is a frequency-dependent and polymorphic version of (8) in Barfield et al. (2011). Importantly, the RHS of equation (C.3) still depends on the class-specific moments 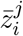 and 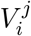. However, after relaxation on the RV space, the quantities 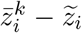 and 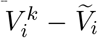 will be *O*(*ε*^2^) and *O*(*ε*^4^) respectively under the oligomorphic approximation (see Online Appendix S.3). Hence, to leading order, we can replace the class-specific morph means and variances by the corresponding moments of the RV-weighted distribution. This substitution thus introduces a small error, but it will be quantitatively acceptable as long as the morph variances remain small. This will notably be the case near evolutionary endpoints under stabilising selection, but our simulations show that the approximation is also accurate away from evolutionarily singularities. With this last approximation, we obtain equation (29) in the main text.

We can also calculate the dynamics of the morph variances, either using a quasi-equilibrium approach (as shown in Online Appendix S.6 for a two-class model), or by calculating the dynamics of the RV-weighted morph variances 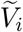. This latter approach yields the folowing Price equation

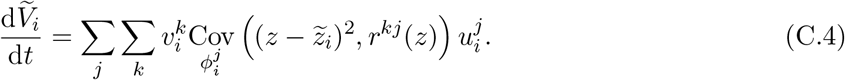

The quadratic term in the covariance can be rewritten using the morph mean 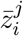 as follows: 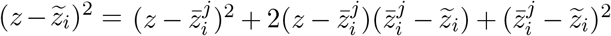. We can use this decomposition to rewrite equation (C.4) as

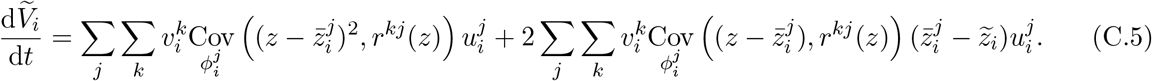

Using the Taylor expansion (6) in the main text to rewrite the covariance terms, we obtain

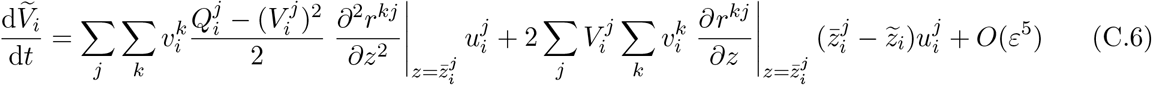

With the Gaussian closure approximation 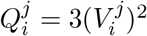 and again using 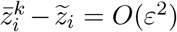 and 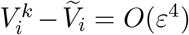, we finally obtain

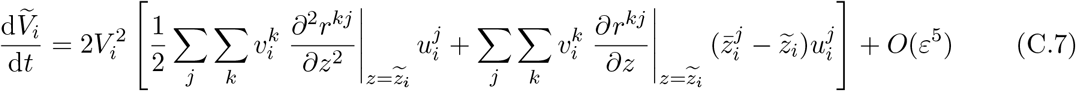

which can be rewritten in matrix form as equation (21) in the main text.

## Appendix D: Two-class models

Two-class models represent a fundamental tool to analyse structured populations. We therefore develop here how our oligomorphic approximation, together with the RV projection, lead to simple dynamical equations in this case.

Consider a population structured in two classes *A* and *B*. As shown in the general case, the dynamics of morph means and variances are given by the following system of equations:

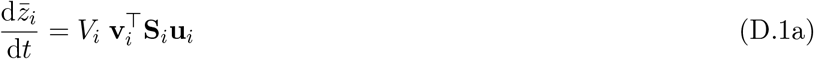

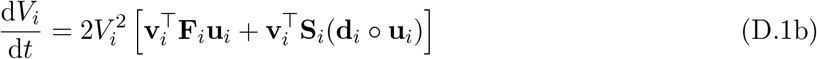

When there are only two class, it is straightforward to show that

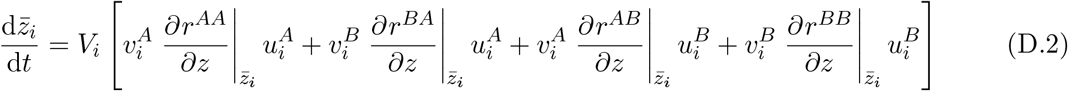

where the class-reproductive values satisfy the following quasi-equilibrium relationship:

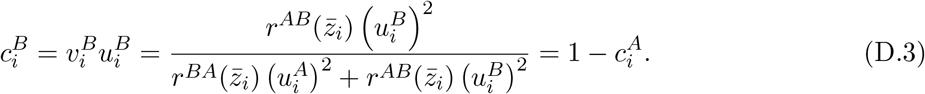

which can be derived from the equations 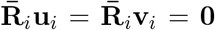 along with the normalisation condition 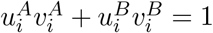 (see Online Appendix S.4).

For the variance dynamics, we have similarly

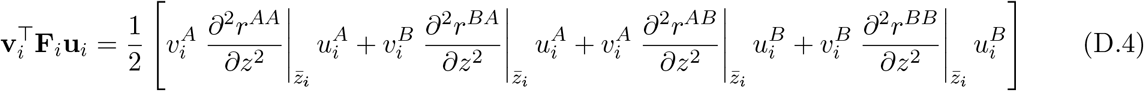

and for the second term in equation (D.1b), which we note 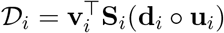, we have

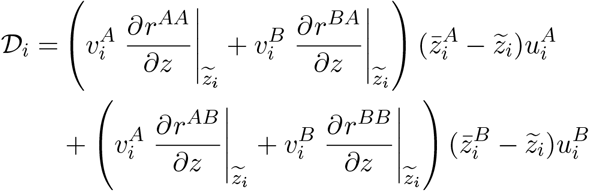

Because 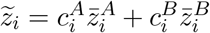, we have 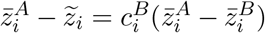 and 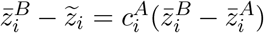, and therefore

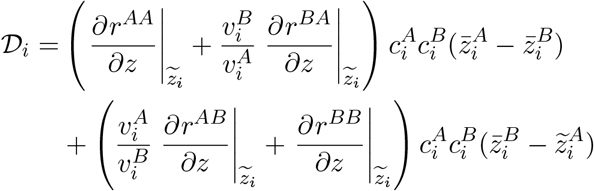

where we have used the definition 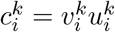. This leads to

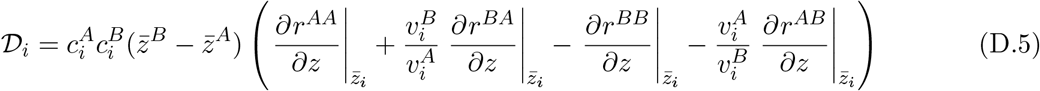

More progress can be obtained if we treat the difference 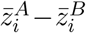 as a fast variable (see Online Appendix S.2 and Lion (2018b)). In a two-class model, we have

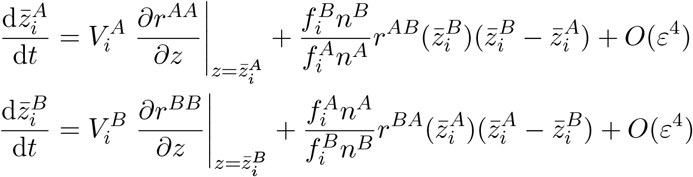

Using these two equations to calculate the dynamics of 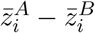 and setting the RHS of the resulting equation to zero leads to the following quasi-equilibrium approximation:

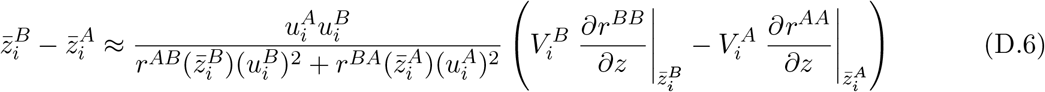

which can be rewritten using equation (D.3) as

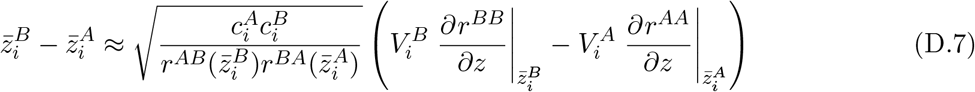

We can then use the morph-level closure approximation 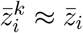 and 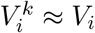 to obtain finally

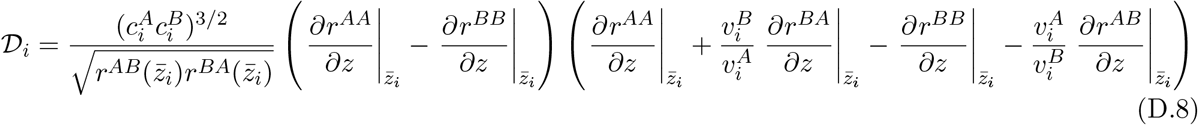

Interestingly, equation (D.8) shows that directional selection will have a significant effect on the dynamics of variance if three conditions are met: (1) there is enough differentiation between the two classes, as measured by the product of class reproductive values 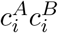, (2) the slopes of the functions *r^AA^* and *r^BB^* at the morph means are sufficiently different, and (3) the marginal reproductive outputs of *A* and *B* individuals are sufficiently different. The latter condition is satisfied when the second bracketed term is non-zero. Note that the ratios 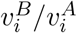 and 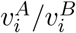 can be interpreted as conversion factors to evaluate the *A* and *B* descendants in the same currency.

Finally, we note that, when 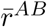 and 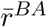 are independent of the trait *z*, as in our migration-selection models, equation (D.8) can be simplified as

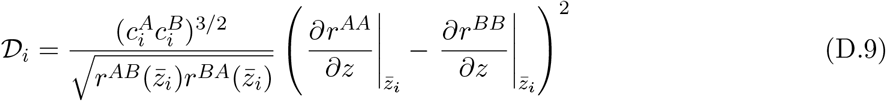

## Appendix E: Mutation

The impact of mutation on the oligomorphic dynamics will depend on the specific mutation model one chooses. For simplicity, we assume here we assume that mutation occurs independently of reproduction, at rate *μ*, and that the mutation effects follow a distribution *M_k_* with mean 0 (mutation has no directional effect) and variance 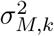. Thus, mutation from phenotype *y* to phenotype *z* is determined by a mutation kernel *m^k^*(*z*, *y*) in class *k*, such that, for all *y*, *∫ m^k^*(*z*, *y*)d*z* = 1, *∫ zm^k^*(*z*, *y*)d*z* = *y* (unbiased mutation) and 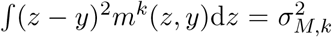. With these assumptions, the dynamics of the density of morph *i* individuals, 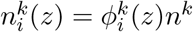 can be modified as follows:

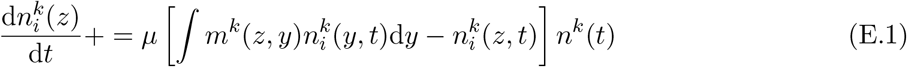

where the + = notation means that we just add the term on the right-hand-side to the results in absence of mutation. Note that equation (E.1) assumes that mutation does not allow transitions across morphs, which will be satisfied if mutation is sufficiently local and the morphs sufficiently distinct. It is easy to see that, if we integrate equation (E.1) over *z*, the mutation term vanishes, and therefore mutation has no effect on the dynamics of the densities 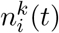.

From equation (E.1), it is straightforward to derive

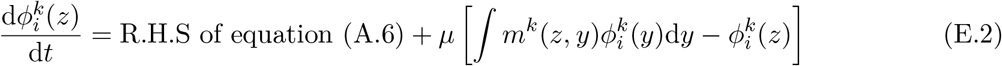

and following the steps in equation A, we can also show that mutation does not affect the dynamics of morph frequencies.

Multiplying equation (E.2) by *z* and integrating, we obtain

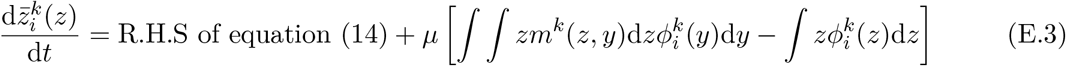

Assuming that mutation is unbiased, so that *y* = *∫ zm^k^*(*z*, *y*)d*z*, it follows that the mutation term in equation (E.3) vanishes, so that mutation has no impact on the dynamics of morph means.

To calculate the variance dynamics, we multiply (E.2) by 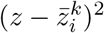 and integrate to obtain

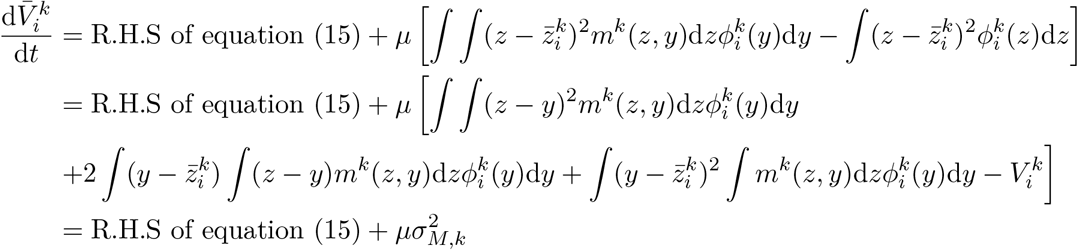

Hence, mutation adds a term 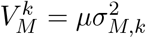 to the dynamics of morph variances. *V_M,k_* is the mutational variance in class *k* (Kimura, 1965; Lande, 1975; Sasaki & Dieckmann, 2011). Finally, to derive the dynamics of the RV-weighted morph variance 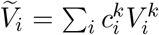, we simply need to add a term 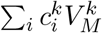 to the right-hand-side of equation (C.7), which yields equations (20) and (21) in the main text.

## Appendix S: Online Appendix for “Multi-morph eco-evolutionary dynamics in structured populations”

### S.1 *Justification of equality* (A.4)

To justify equation (A.4), we first start from the natural assumption that individuals with the same phenotypes have the same per-capita growth rate in each class (say *k*) whichever morph they belong:

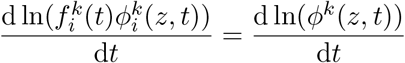

This is equation (A.2) in Appendix A. We write the same relationship in class *j*:

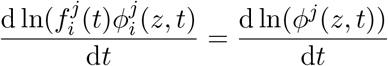

and subtracting these two equations yields

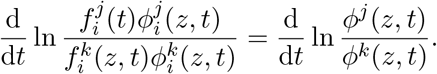

Integrating both sides over *t* yields

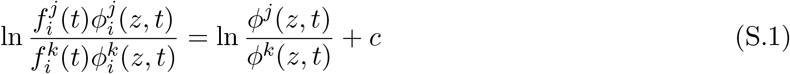

where

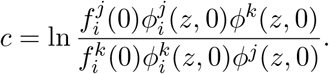

Hence,

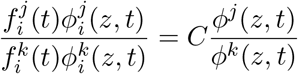

where

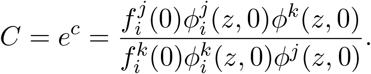

Therefore, if

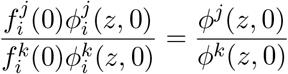

initially holds, then

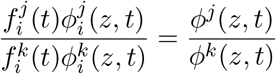

holds for *t* ≥ 0. This justifies equality (A.4).

### S.2 Projection on RV space: quasi-equilibrium approach

The goal of this appendix is to show how some results and arguments derived in Lion (2018b) for unimodal distributions can be extended to multimodal distributions under the oligomorphic approximation by working at the morph level. Let us define **u**_*i*_ the vector with elements 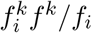, **U**_*i*_ = diag(**u**_*i*_), **C**_*i*_ the matrix with elements 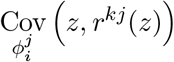, and **d**_*i*_ the vector of morph-specific scaled phenotypic differentiation (with elements 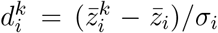 where 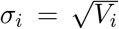 is the standard deviation of the distribution of morph *i*). We can then write equation (C.1) as

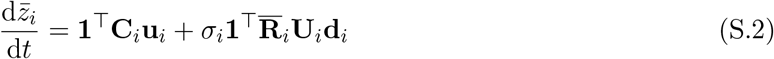

which has the same form as equation (15) in Lion (2018b), but is morph-specific. Similarly, the dynamics of **d**_*i*_ can be put in the form:

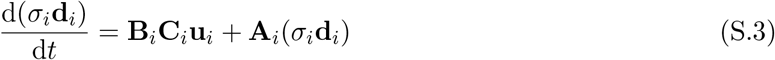

which has the same form as equation (A2) in Lion (2018b). Hence, if we assume, as in Lion (2018b), that the unimodal morph distributions are tightly clustered around the mean (which is the crux of the oligomorphic approximation), we can follow the same approach as in that paper, and derive a quasi-equilibrium approximation for **d**_*i*_. This eventually yields:

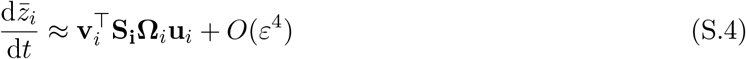

where 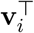 and **u**_*i*_ are calculated as the left and right eigenvectors of **R**_*i*_ associated to eigenvalue 0, keeping only the *O*(1) terms, 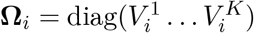 and the matrix **S**_*i*_ has elements *∂r^kj^*/*∂z* evaluated at 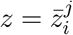. To leading order, we can replace 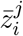 and 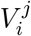 by 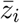 and *V_i_* (because both 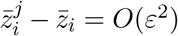 and 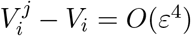 on the slow time scale, as shown in Online Appendix S.3) to obtain equation (19) in the main text. Doing so only contributes an *O*(*ε*^4^) error term.

Finally, a similar reasoning can be applied to the dynamics of variance, and we conjecture that the dynamics of all moments of the distribution *ϕ_i_*(*z*, *t*) can be approximated by those of 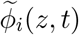 for small *ε*.

### S.3 Separation of time scales

In this appendix, we first show that the phenotypic differentiations 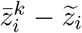 converge to zero on the fast time scale, so that we can assume 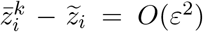 on the fast time scale. We then use a perturbation expansion approach to prove the separation of time scales between class densities and morph frequencies on the one hand, and morph means on the other hand. The arguments for the dynamics of variance are similar.

#### Dynamics of phenotypic differentiation on the fast time scale

Let us introduce the deviations 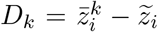, which measure the difference between the mean trait in class *k* and the RV-weighted mean trait 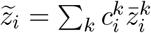. We then have

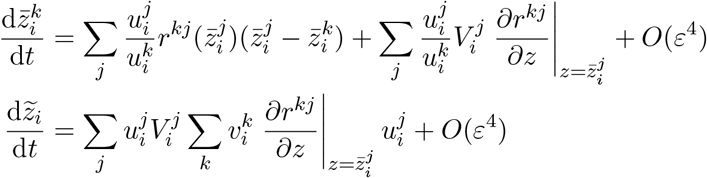

Subtracting the two equations and keeping only *O*(1) terms yields

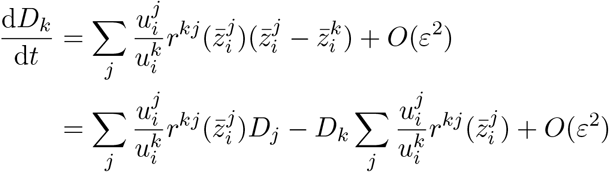

Writing the differentiation as the perturbation expansion 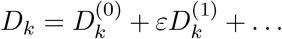, it follows that the zeroth-order term satisfies

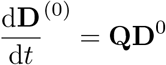

where the matrix **Q** has the following elements

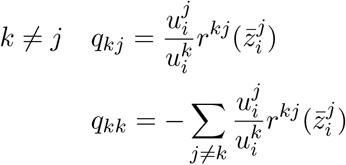

Because 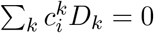 by definition, the system is overdetermined, but we can remove one redundant equation and write the dynamics of **D**_*_ = (*D*_1_… *D*_*K*–1_). This gives

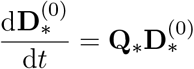

where the matrix **Q**_*_ is a (*K* – 1) × (*K* – 1) matrix obtained from **Q** by expressing *D_K_* as a function of the elements of **D**_*_ (Lion, 2018b). If we assume for convenience that the phenotypic differentiation does not blow up and the system of class densities, morph frequencies and phenotypic differentiations reaches an equilibrium in the limit *ε* → 0, and **Q**_*_ is invertible at this equilibrium, then we can characterise the equilibrium as **D**^(0)^ = **0**. Note that the same reasoning allows us to show that **D**^(1)^ = 0, and therefore

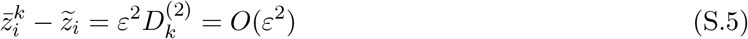

This justifies that we can replace 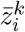 by 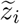 on the slow time scale (i.e. after relaxation of the fast dynamics). Note that the argument above implicity requires that the classes must not be isolated, otherwise the transition rates *r^kj^* are all zero for *j* ≠ *k*, and there is no homogeneisation of the morph means on the fast time scale.

For the variance dynamics, plugging (S.5) into equation (15) shows that the leading-order term of the dynamics of the variance differentiation 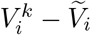 is

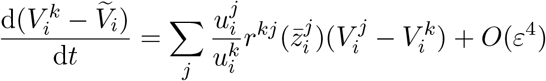

which has the same form as the dynamics of **D**^(0)^ and therefore the same argument shows that

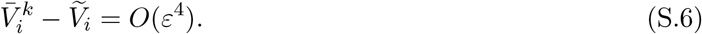

#### Two-class case

For the two-class case, we can use milder assumptions. Let 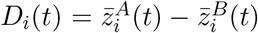 and *L_i_*(*t*) = *D_i_*(*t*)^2^. Then, if we simply assume that *r^AB^*(*z*) > 0 and *r^BA^*(*z*) > 0 for any *z* (which will be realistic for many ODE ecological models), we have in our *O*(1) system

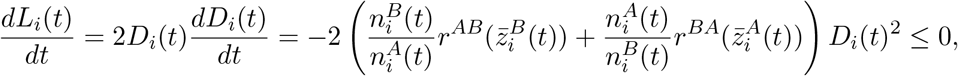

where the equality holds only when *D_i_* = 0. Therefore 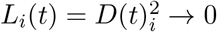.

#### Separation of time scales: perturbation expansion

Here, we use a perturbation expansion to analyse the separation of time scales in the oligormophic equations. The argument is valid for a given morph *i*, so for simplicity we drop the *i* subscript. We have for the dynamics of class densities

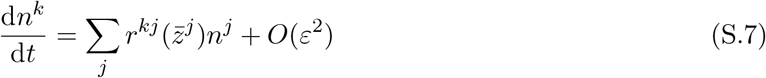

and for the dynamics of class means

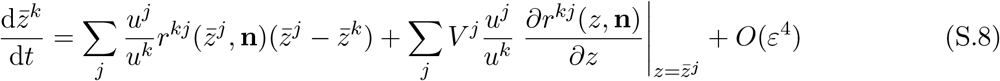

It is clear that the leading-order term of equation (S.7) is *O*(1). But so is the leading-order term of equation (S.8). However, we will see that after relaxation of the fast dynamics, equation (S.8) is *O*(*ε*^2^).

To show this, we use the following perturbation expansion:

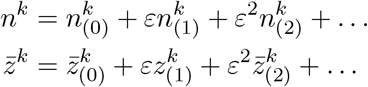

We first solve the system for the zeroth-order terms:

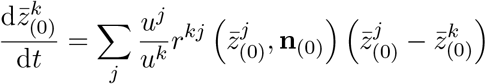

This system is obtained by Taylor-expanding equation (S.8) and keeping only *O*(1) terms. These dynamics tend to homogeneise the values of the mean traits, so (as we’ve just shown above) we have at equilibrium

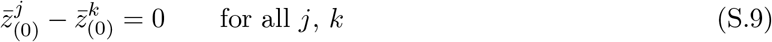

We can use this quasi-equilibrium result to obtain the following system for the dynamics of **z***k*_(1)_

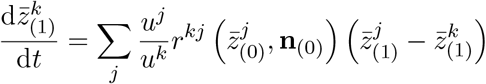

This system is obtained by Taylor-expanding equation (S.8) and keeping only *O*(*ε*) terms. Again, this yields at equilibrium

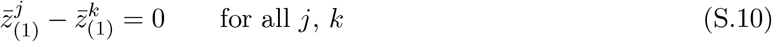

In particular, this means that

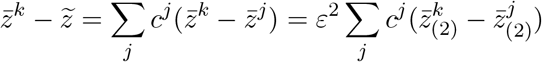

and therefore

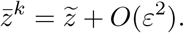

We then use the quasi-equilibrium solutions of (S.9) and (S.10) to derive the dynamics of the *O*(*ε*^2^) terms as

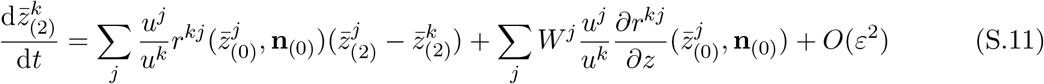

where *V^j^* = *ϵ*^2^*W^j^*.

Thus, after relaxation of the fast dynamics, we have

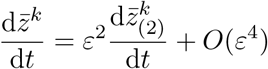

where 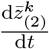 can be calculated using equation (S.11).

Note that equation (S.11) depends on the zeroth-order terms of densities **n**, but actually, if we use **n** instead of **n**_(0)_ in equation (S.11), the error we make will be absorbed in the *O*(*ε*^2^) remainder. Similarly, it is possible to replace 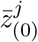 by 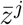 or 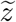 in the arguments of *r^kj^*, so we can also write more simply

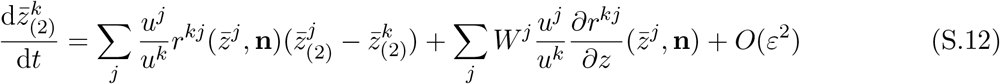

In words, this analysis indicates that, starting from distinct morph means in each class, the morph means first change quickly to become clustered around an average value (i.e. 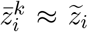) then change more slowly along the slow manifold. On this manifold, the densities are well approximated by the quasi-equilibrium solution of **n**_(0)_.

Note that the dynamics of the RV-weighted mean trait, 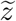, is *O*(*ε*^2^) from the start, which is why it is much easier to use it to describe the slow dynamics.

### S.4 Quasi-equilibrium approximation of reproductive values in two-class models

In two class models, the equations 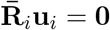 and 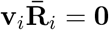 can be rewritten as

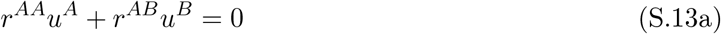

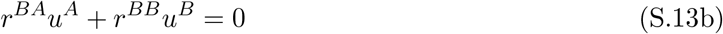

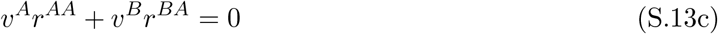

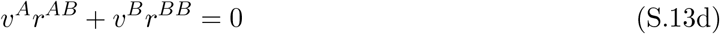

where for simplificity we remove the dependency on the morph (i.e. we write 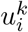 as *u^k^*, 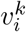 as *v^k^*, and 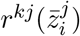 as *r^kj^*.

Using equation (S.13c), we can rewrite the normalisation conditions *u^A^v^A^* + *u^B^v^B^* = 1 as

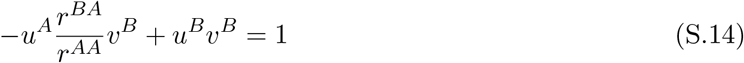

which leads to

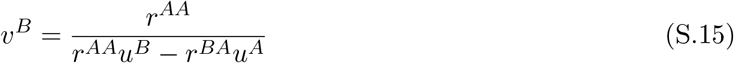

and, using equation (S.13a), which can be rewritten as *r^AA^* = −*r^AB^u^B^/u^A^*, we obtain finally

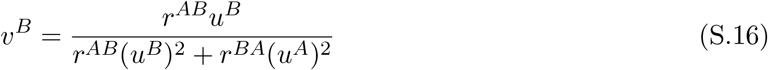

A similar equation can be obtained for *v^A^* (we just need to swap the A and B superscripts) and multiplying by *u^B^* yields

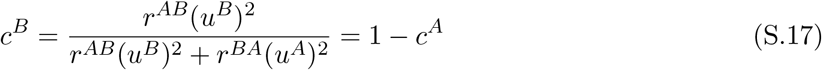

which is equation (D.3).

### S.5 Oligomorphic dynamics in a two-class migration-selection model

We use equation (22) in the main text to derive the oligomorphic dynamics when there are only two classes and when the only transitions between distinct classes correspond to migration and are independent of the focal trait under consideration. That is, we have:

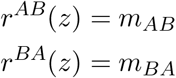

With this simplifying assumption, we obtain the following equations, which will be used to analyse our three Examples:

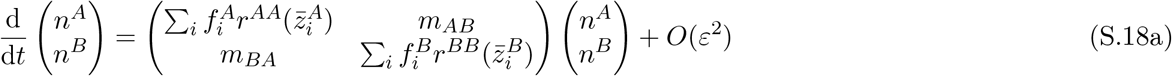

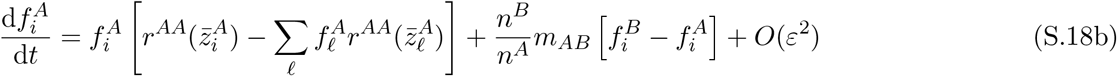

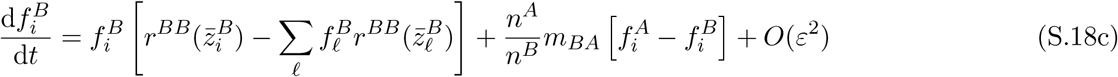

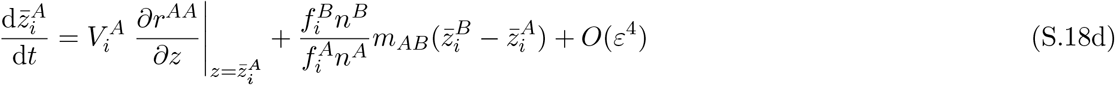

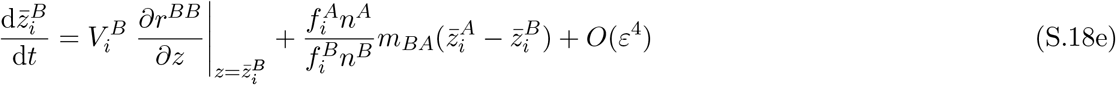

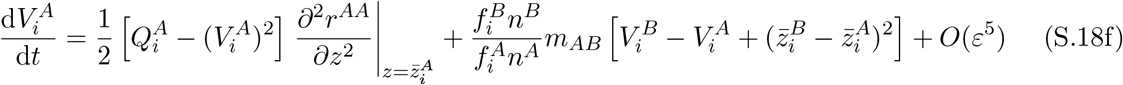

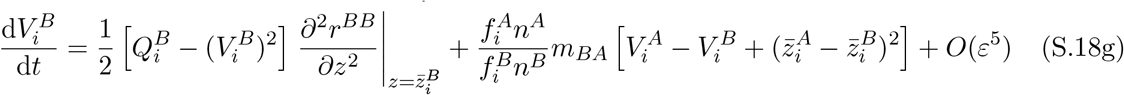

### S.6 Example 1: A two-habitat local adaptation model

In this appendix, we carry out an explicit analysis of a specific two-habitat model to revisit the results of Débarre et al. (2013) (see also Ronce & Kirkpatrick (2001) and Mirrahimi & Gandon (2020)). As in that paper and in Online Appendix S.5, we consider a population of individuals distributed over two habitats, *A* and *B*, coupled by migration. Each habitat is characterised by a habitat-specific optimum (*θ_A_* and *θ_B_*, respectively). The transition rates between classes are then

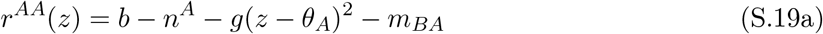

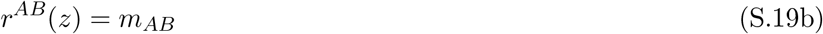

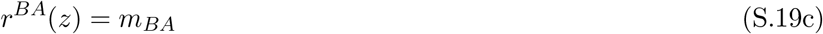

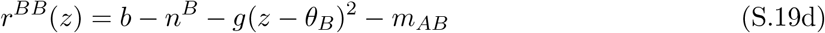

where *b* is the fecundity rate and *g* is the fecundity cost. We use quadratic cost functions for simplicity, so that the cost is minimal at the habitat’s optimum. Also, in contrast to Débarre et al. (2013), we consider asymmetric migration rates, with *m_jk_* the migration rate from habitat *k* to habitat *j* (see also Mirrahimi & Gandon (2020)). Note that we assume migration rates do not depend on the focal trait, which will lead to simplifications as the partial derivatives of *r^kj^*(*z*) will vanish for *j* ≠ *k*.

Note that this model can be simply obtained from the competition model in Online Appendix S.7 by assuming *a*(·) = 1.

#### S.6.1 Oligomorphic approximation

To derive our oligormorphic approximation for this model, we combine equations (S.18) with the following relationships

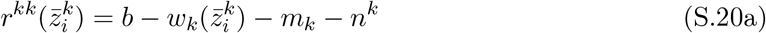

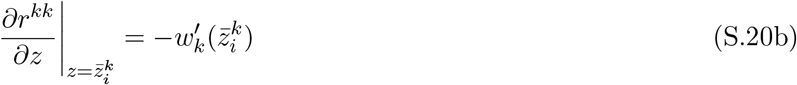

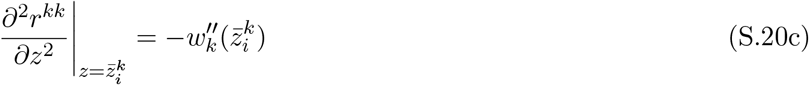

where *w_A_*(*z*) = *g*(*z* – *θ_A_*)^2^, *w_B_*(*z*) = *g*(*z* – *θ_B_*)^2^, *m_A_* = *m_BA_* and *m_B_* = *m_AB_*.

##### Reproductive values

The resulting system can be numerically solved, but we can get some further simplifications using reproductive values. Using equation (D.3), we see that class reproductive values satisfy at quasi-equilibrium

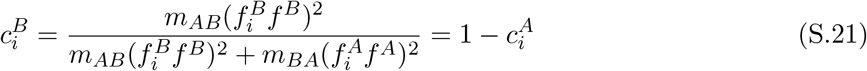

where the morph and class frequencies are calculated using the *O*(1) terms of equations (S.18a)–(S.18c).

##### Morph means

Using equations (S.4) and (C.2), it is straightforward to derive the following equation for the dynamics of the morph mean, 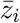, and the RV-weighted morph mean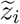. We obtain

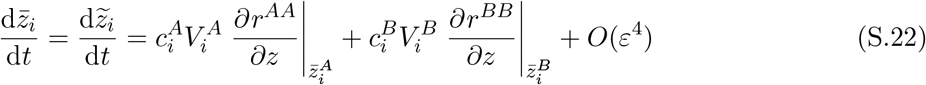

If we only want to keep *O*(*ε*^2^) terms, it is sufficient to replace 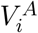 and 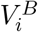 by *V_i_*, the morph variance, and 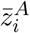 and 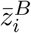 by the morph mean 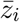 (Online Appendix S.3). We then obtain for our model:

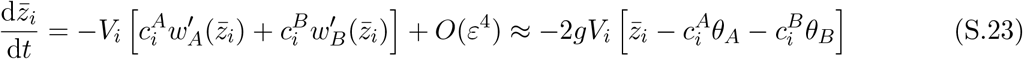

An explicit derivation of equation (S.22) can also be obtained by calculating the dynamics of 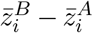 which is simply obtained by substracting equations (S.18d)–(S.18e). Then, treating this phenotypic differentiation as a fast variable (Lion (2018b); Online Appendices S.2; S.3), we can set the right-hand side of the resulting equation to zero, and solve for 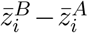. This yields the following quasi-equilibrium approximation

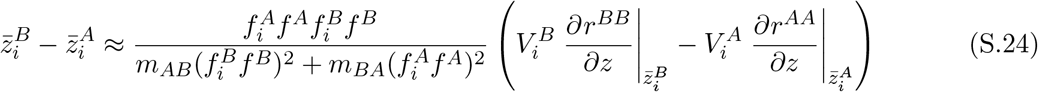

which can be written using equation (S.21) as

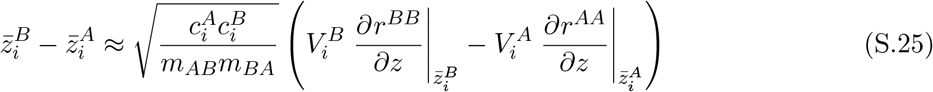

Plugging the resulting expression into equations (S.18d) and (S.18e) yields

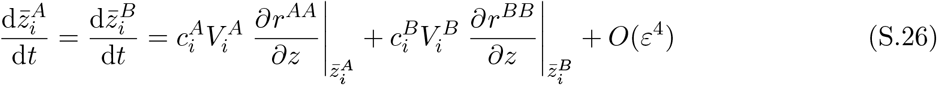

which entails, assuming that the morph and class frequencies are calculated on their quasi-equilibrium manifold,

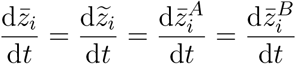

Equation (S.25) shows that 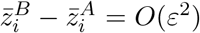, so that 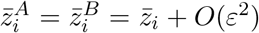. This is consistent with the general argument in Online Appendix S.3 and justifies that we can replace the class-specific morph means by the morph mean in equation (S.22). We then obtain equation (29) in the main text. Figure S.1 shows that the differentiation between morph means 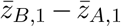 quickly converges to a small value which is well predicted by the quasi-equilibrium approximation.

**Figure S.1:**
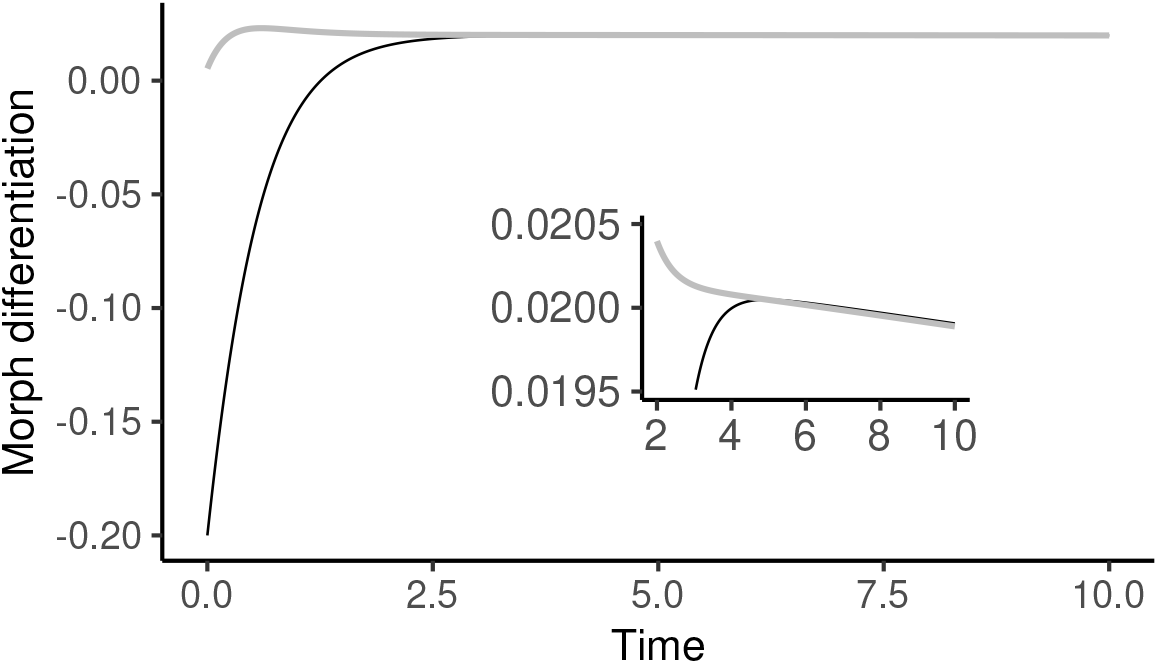
llustration of the relaxation to RV space in a two-class model. The simulation is the same as in figure 2 and figure S.3A, to which the reader is referred for additional details. The morph differentiation 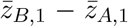 (black) is shown to converge towards the value predicted by the quasi-equilibrium approximation (S.24) (grey line). The inset shows a close-up of the dynamics.

##### Morph variances

Similarly, we can derive the dynamics of the difference in morph variances from equations (S.18f)–(S.18g). With Gaussian closure approximation (such that 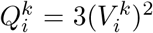) and using equation (S.21), we obtain after some rearrangements the following quasi-equilibrium approximation:

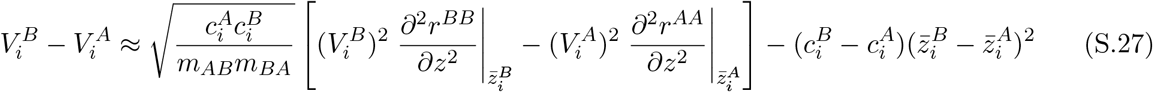

Plugging this into equations (S.18f) and (S.18g) yields, again after some rearrangements

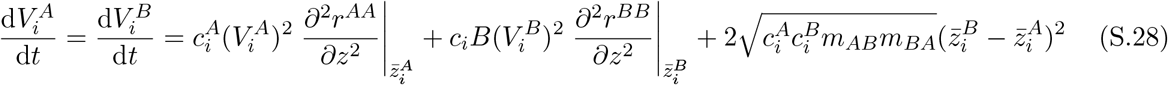

and using equation (S.25) finally yields

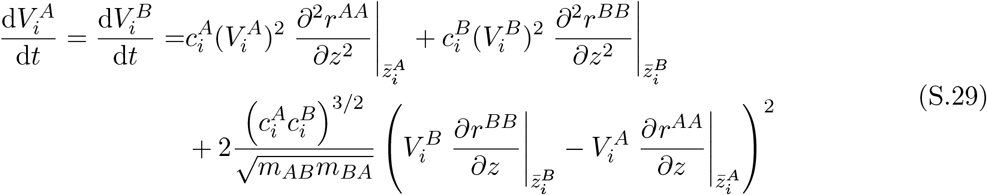

which again implies

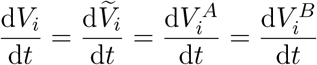

Another route to this result is to start with the dynamics of the RV-weighted variance, given by equation (21). This gives for our model

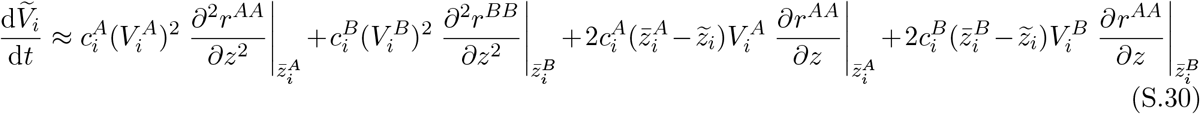

(recall that 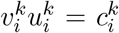). Noting that 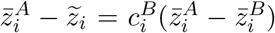 and 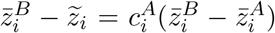, and using equation (S.25) finally yields

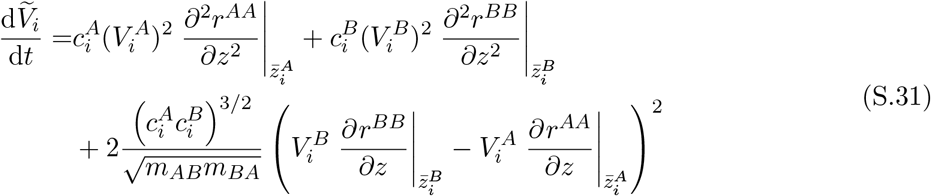

From equation (S.27) we see that 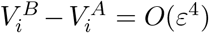, so that 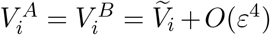. The errors made by approximating the class-specific variances by 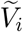 in equation (S.31) will therefore be of higher-order than the leading-order terms.

#### S.6.2 Quadratic functions

For the model with quadratic functions, equations (S.22) and (S.31) simplify to

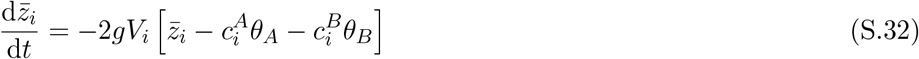

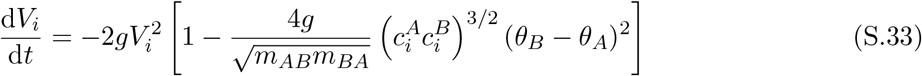

Equation (S.32) shows that at equilibrium morph means are equal to the reproductive-value weighted average of habitat optima, 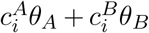. As shown in Sasaki & Dieckmann (2011), the corresponding equilibra are all evolutionarily stable if and only if d*V_i_*/d*t* < 0 for all morphs, which is equivalent to

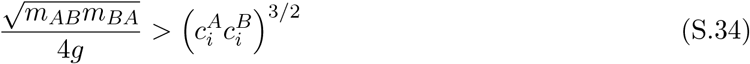

where we have set *θ_A_* = 0 = 1 – *θ_B_* without loss of generality.

#### S.6.3 Solutions

We consider that the population is composed of two morphs (1 and 2). From equations (S.18b) and (S.18c), together with equations (S.19), we can thus write the dynamics of the frequencies of morph 1 in habitats *A* and *B* as follows:

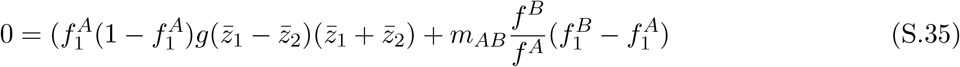

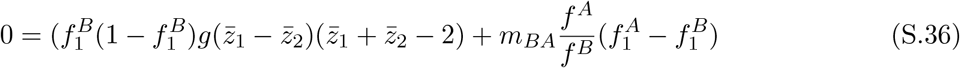

Multiplying the first equation by *f^A^*/(*m_AB_f^B^*) and the second by *f^B^*/(*m_BA_f^A^*), then taking the sum, gives

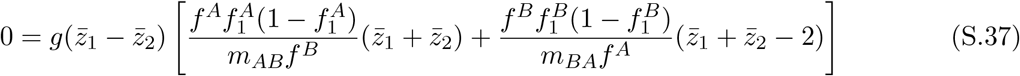

This is satisfied either if 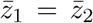 which corresponds to a single-morph equilibrium, or if the term between brackets is zero, which leads to a dimorphic equilibrium

##### Dimorphic equilibrium

We start with the dimorphic case, which is simpler to analyse. Setting the term between brackets in equation (S.37) to zero yields

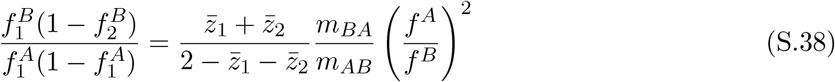

From the equilibrium of equation (S.32), we have 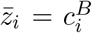, and because 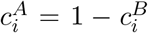, we thus have 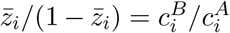. From equation (S.21), we then have 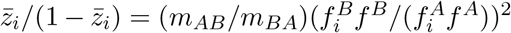, which we can use to simplify equation (S.38) as

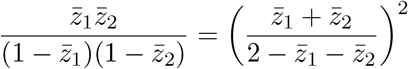

which can be rearranged as

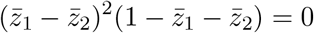

Hence, because 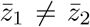 for the dimorphic equilibrium, the morph means must satisfy 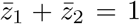. Plugging this condition into the dynamics of morph frequencies, then solving for 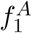 and 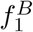 and calculating 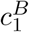 yields

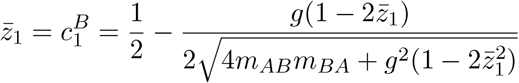

Solving for 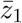 finally yields, for 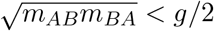

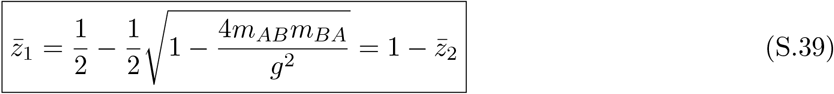

which, using equation (S.33), is evolutionarily stable if 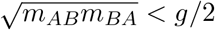. Equation (S.39) corresponds to the results of Mirrahimi & Gandon (2020) and, for the symmetric migration case, to those of Débarre et al. (2013).

Equation (S.39) can be used to calculate the morph frequencies (see companion Mathematica notebook). In the same notebook, we also derive the following expressions for the equilibrium densities

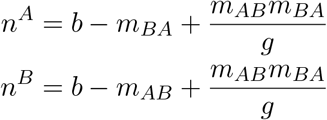

For symmetric migration, we recover the results in Table 1 of Débarre et al. (2013).

Because the dimorphic equilibrium is characterised by the two morphs having different frequencies in the two habitats, the habitat-specific trait distributions are distinct. We can characterise these equilibrium distributions by calculating their moments (see companion notebook):

- the mean trait in habitats *A* and *B* (and the differentiation 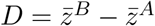)

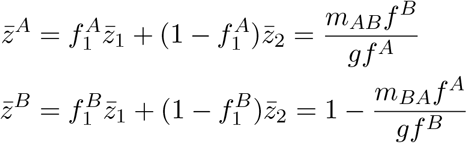
- the global mean trait

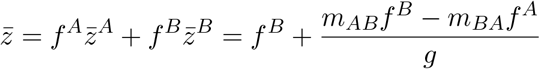

where the first term is the mean of the two optima (*f^A^θ_A_* + *f^B^θ_B_* = *f^B^*) and the second term is the deviation caused by the migration-selection balance. For symmetric migration, we have 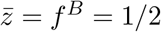.
- the variance in habitat *A* in the absence of mutation-selection balance (i.e. assuming 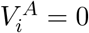 at equilibrium)

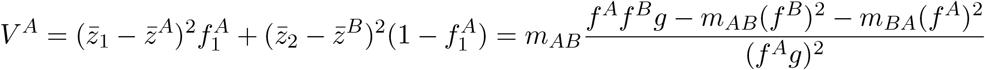
- the third moment in habitat *A* in the absence of mutation-selection balance (i.e. assuming 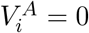 at equilibrium), assuming the morph distribution is not skewed (e.g. 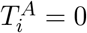)

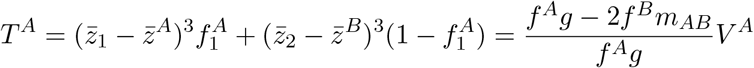

Note that, for symmetric migration, we recover the results of Débarre et al. (2013) (column 2 in their Table 1). In figure S.2, the dynamics of the moments of the trait distributions in habitats A and B are presented and compared with the analytical predictions.

**Figure S.2:**
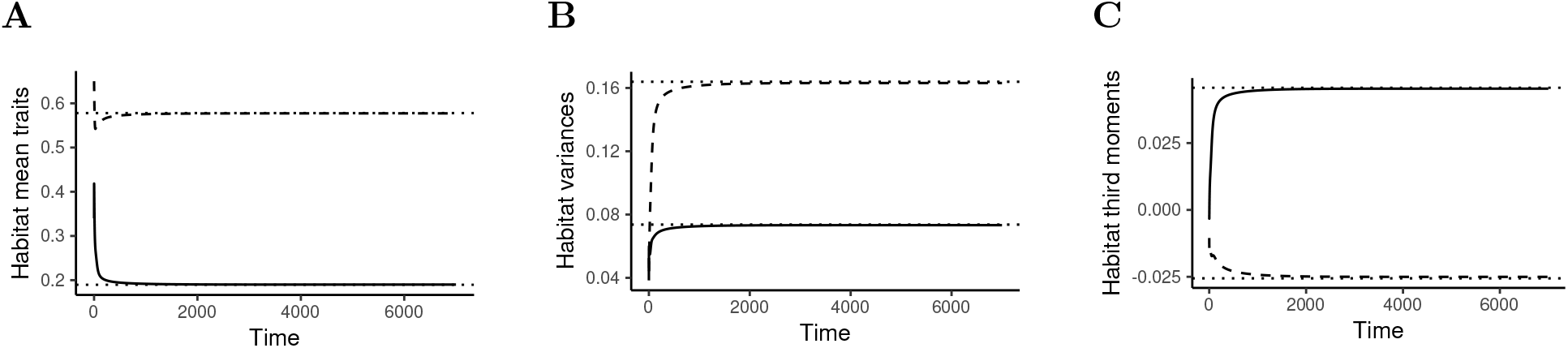
Dynamics of habitat-specific moments. The means (a), variances (b) and third moments (c) of the trait distributions in habitat A (solid lines) and B (dashed lines) and shown to converge to the values predicted by the analytical formulae (dotted horizontal lines). Parameters as in figure 3B in the main text.

##### Local adaptation

At the bimodal equilibrium, the two morphs have different frequencies in the two habitats and therefore the habitat-specific distributions are distinct (figure 3B). As shown by Débarre et al. (2013), local adaptation in this model is proportional to the level of habitat differentiation in mean traits, which can be calculated as

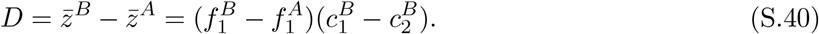

This shows that, at equilibrium, local adaptation depends (1) on the difference in morph frequencies between the two habitats, and (2) on the difference between the class reproductive values of the two morphs. Using the above results, habitat differentiation can be calculated as:

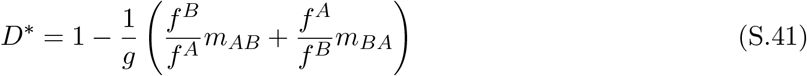

which simplifies to *D*\* = 1 – 2*m/g* for symmetric migration, as found in Débarre et al. (2013). Higher migration thus leads to lower local adaptation.

##### Single-morph equilibria

In the single-morph case, we have only one morph with frequencies 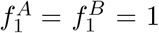. The equilibrium densities and morph mean can be calculated from the following system of equations:

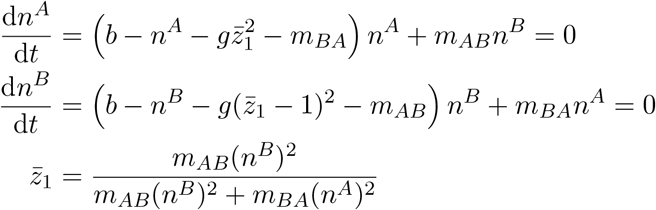

where the latter equation simply states that the mean trait is equal to the class reproductive value 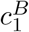.

The system can only be fully solved numerically, except for symmetric migration where at least one solution (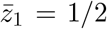 and *n^A^* = *n^B^* = *b* – *g*/4) can be analytically calculated. Depending on the region of parameter space (and in particular the values of the migration rates) there is typically either one or three solutions of the system.

Note that, in the limit where *m_AB_* = *m_BA_* = *m*, we have 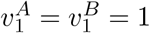 and therefore 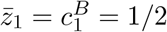. The “symmetric monomorphic” singularity found by Débarre et al. (2013) thus corresponds to the case where both habitats have equal reproductive values. As found by Débarre et al. (2013), this solution is evolutionarily stable if *m* > *g*/2, which can be checked using condition (S.33).

##### Bistability

For some parameters values, the system can exhibit several evolutionary attractors (i.e. convergent and evolutionarily stable points, Geritz et al. (1998)), notably the dimorphic equilibrium and one or two single-morph equilibria. The endpoint of the eco-evolutionary dynamics is then determined by the initial conditions. This bistability is illustrated in figure S.3 for a specific example, and the full bifurcation diagrams of the model for *m_AB_* = 0.8 are shown in figures S.4. Note however that, as already found by Débarre et al. (2013), the basin of attraction of the unimodal equilibrium is relatively narrow so that a little mutation is sufficient to push the dynamics towards the bimodal equilibrium. This explains why the simulations of the full model (the black dots in figures S.4A and S.4B) typically converge towards the bimodal equilibrium when it exists. Thus, while the oligomorphic analysis predicts bistability when *m* < *m_c_*, with some initial conditions leading to unimodal equilibrium distributions, a global stability analysis shows that the bimodal distribution is the more robust evolutionary outcome (Mirrahimi & Gandon, 2020).

**Figure S.3:**
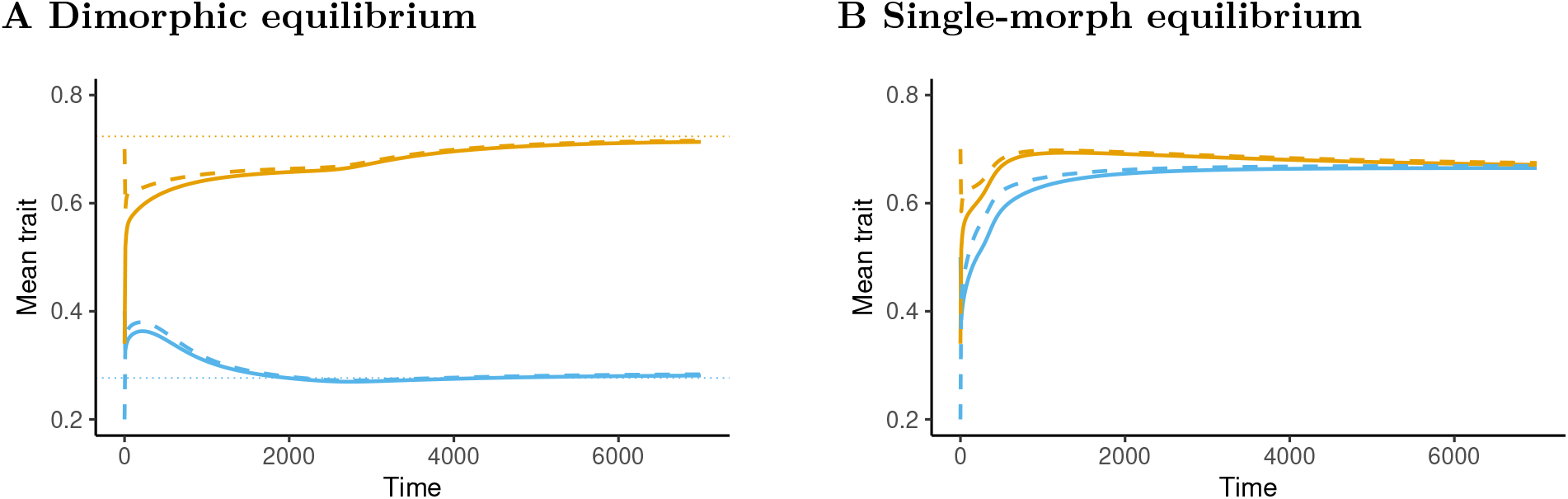
Illustration of the bistable dynamics of the model for sufficiently low values of the geometric mean of the migration rate, 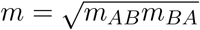. The dynamics of the mean trait of morph 1 (blue) and 2 (black) in habitat A (solid lines) and B (dashed lines) are shown, either leading to a polymorphic equilibrium (left panel) or to a monomorphic equilibrium (panel b). The only difference between the two simulations is the initial trait value of the first morph in habitat A, which is 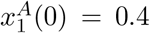 in panel (a), and 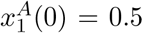 in panel (b). Other initial conditions: *n_A_*(0) = *n_B_*(0) = 1, 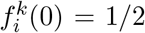 (for *i* = 1, 2 and *k* = *A*, *B*), 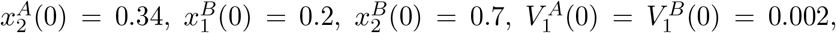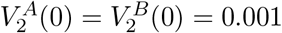. Parameters: *m_AB_* = 0.8, *m_BA_* = 1, *g* = 2, *b* = 1.

#### S.6.4 Accuracy of the RV projection

How accurate is it to replace the equations (22) the projection of RV space ? Figure S.4 shows that the quantitative match is very good, except in a small region of parameter space between *m_AB_* = 0.8 and *m_BA_* = 1, where the RV projection does not accurately predicts the single-morph solution. This corresponds to a point where migration is close to symmetric and the single-morph solution actually becomes a repellor. Thus the dynamics converge towards a point where selection is disruptive but a dimorphism cannot persist. As shown in figure S.5B, this causes the build-up of subtantial differentiation between the morph means in habitats *A* and *B*, at which point the morph-centred reproductive-value-weighted oligomorphic approximation breaks down.

#### S.6.5 Effect of mutation

In this section, we give additional results for the analysis of the effect of mutation variance shown in figure 4B. In these simulations, we assume 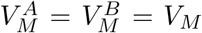. Figure S.6 shows how the mutational variance affects the equilibrium trait distributions in habitat A (top panel) and B (middle panel), and the morph frequencies (bottom panel). There is a sharp change in behaviour around *V_M_* ≈ 10^-3^, which corresponds to the oligomorphic approximation breaking down when the morph variances become too large and the morph distributions collide: we then shift from a dimorphism with two distinct morphs (distinct frequencies of morph 1 and 2 in each habitat, but the morph means are the same in each habitat 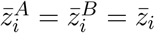) to a case where one morph suddenly goes extinct, but this morph has a distinct mean in each habitat (i.e. 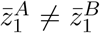). However, the simulations of the full model do not predict this pattern, but rather than the model always converges towards a bimodal distributions with two peaks, albeit with slightly wider variances when *V_M_* is larger. Since the oligomorphic approximation relies on morph variances (i.e. the width of the peaks) being small enough, this is an expected behaviour of our approach. Nonetheless, the oligomorphic approximation remains accurate for relatively large mutational variance, approximately of the same order as the morph variance.

**Figure S.4:**
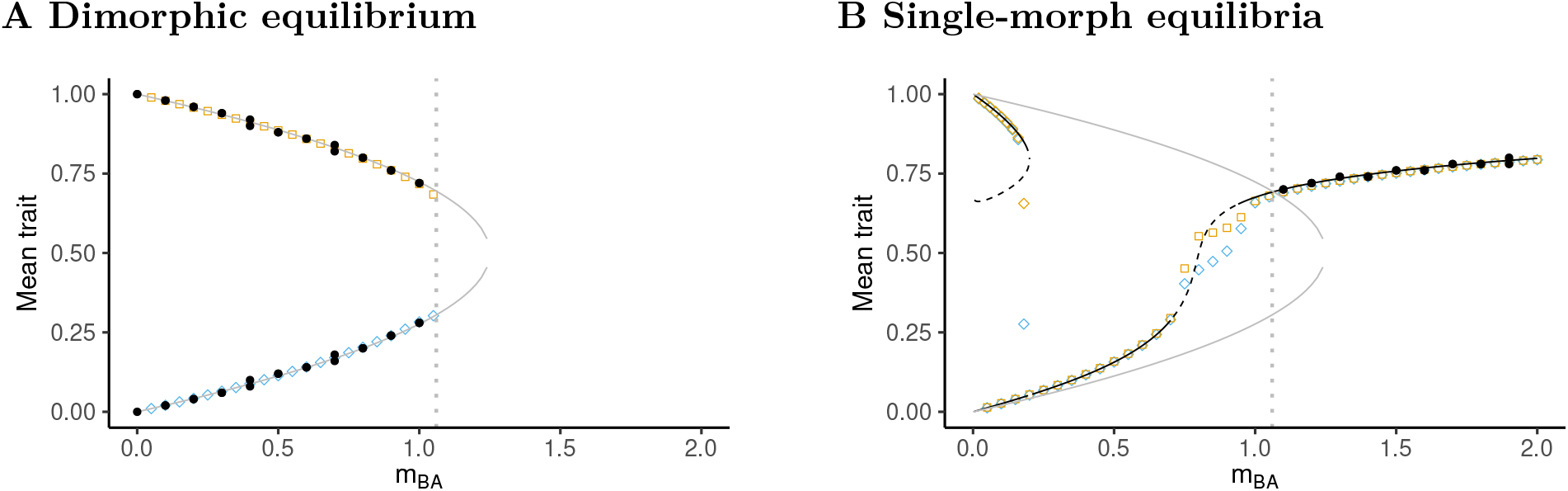
Bistability. Figures (a) and (b) give the bifurcation diagrams for the mean traits. Open shapes give the predictions of a two-morph oligomorphic approximation for the dimorphic (circles, panel b) and single-morph (diamonds, panel c) solutions, for both morphs 1 (blue) and 2 (orange). The gray lines represent the analytical expressions 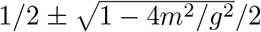, which are shown on both panels (b) and (c) for convenience. The results of the full model, without the oligomorphic approximation, are presented using black dots. On panel (b), the black lines give the predictions of the oligomorphic approximation (solid lines represent evolutionarily stable, and dashed lines evolutionarily unstable solutions). In all panels, the vertical dotted line represents the value *m_BA_* ≈ 1.06 at which the dimorphic equilibrium loses its demographic stability and one of the two morphs goes extinct. Parameter values: *b* = 1, *g* = 2, *m_AB_* = 0.8, *V_M_* = 10^-6^.

**Figure S.5:**
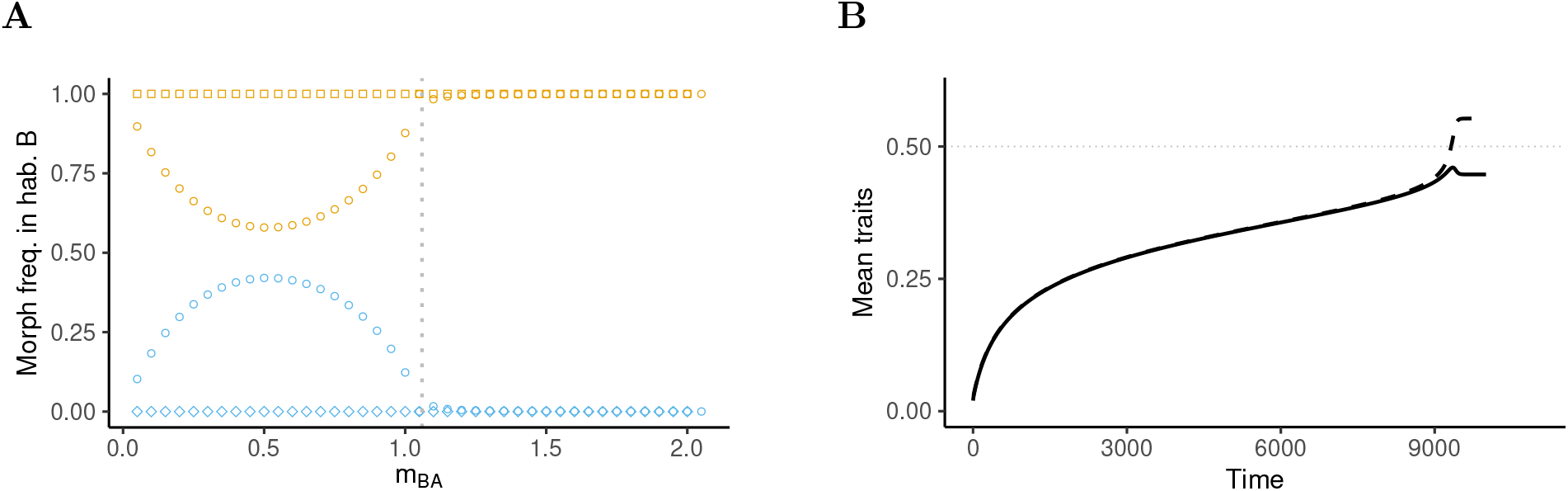
This figure gives some additional results which are helpful to better understand figure S.4 Panel (a) gives the corresponding equilibrium values of the frequencies of morphs 1 (blue) and 2 (orange) for the single-morph (diamonds) and two-morph (circles) equilibria. Panel (b) represents the dynamics of the mean trait in habitat *A* (solid line) and *B* (dashed line) predicted by the single-morph oligomorphic approximation for *m_AB_* = *m_BA_* = 0.8. In all panels, the vertical dotted line represents the value *m_BA_* ≈ 1.06 at which the dimorphic equilibrium loses its demographic stability and one of the two morphs goes extinct. Parameters as in figure 3.

When the mutational variances are different in habitats A and B, we have the following expression for the morph variance at the mutation-selection equilibrium:

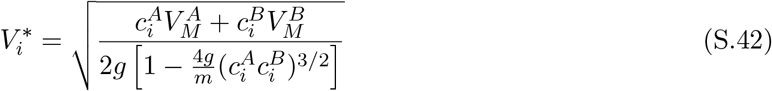

with 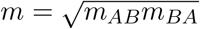. Using the expression (S.39) and the fact that 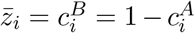 at evolutionary equilibrium, it is straightforward to obtain a closed analytical expression:

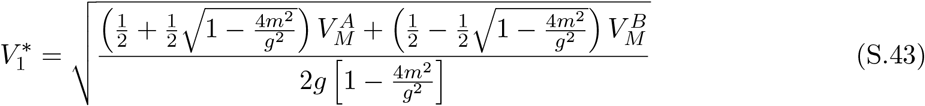

and

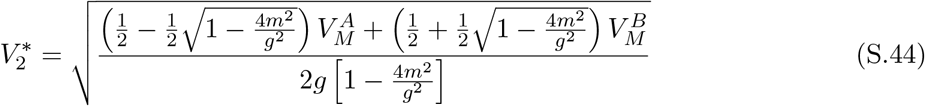

These expressions correspond to the horizontal dotted lines in the top panel of figure 4A.

### S.7 Example 2: A two-habitat resource-competition model

In this appendix, we consider a population of individuals distributed over two habitats, *A* and *B*, coupled by migration. We extend the resource-competition model analysed in Sasaki & Dieckmann (2011) to model migration across classes, and class-specific trait-mediated competition for resources. The transition rates between classes are then

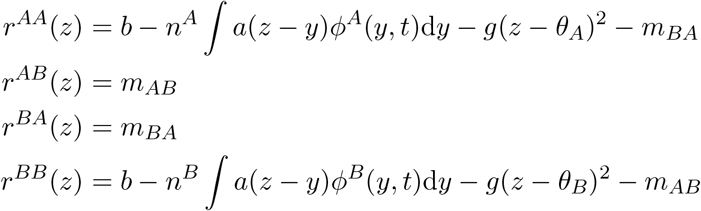

where *b* is the fecundity rate and *g* is a fecundity cost. We use quadratic cost functions for simplicity, so that the cost is minimal at the habitat optima *θ_A_* and *θ_B_*. The function *a*(*z* – *y*) is the competition kernel, which gives the intensity of competition experienced by individuals with trait *z* when they interact with individuals with trait *y*. We assume that the kernel is a symmetric function of the trait difference, so that *a*(*z* – *y*) = *a*(*y* – *z*), and furthermore we assume, as in Sasaki & Dieckmann (2011), that *a*(0) = 1 and *a*′(0) = 0. In the case where only one habitat is present, this corresponds to the model studied by Sasaki & Dieckmann (2011).

#### Oligomorphic dynamics

The dynamics of the densities, frequencies, means and variances are given by system (S.18) We use the oligomorphic approximation to calculate the derivatives of the rates *r^AA^*(*z*) and *r^BB^*(*z*) for the specific case of Example 2. As in Sasaki & Dieckmann (2011), we first Taylor-expand the competition kernel around 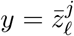 to obtain

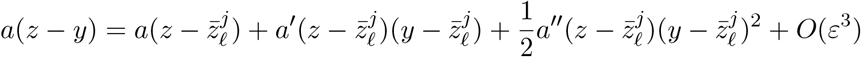

**Figure S.6:**
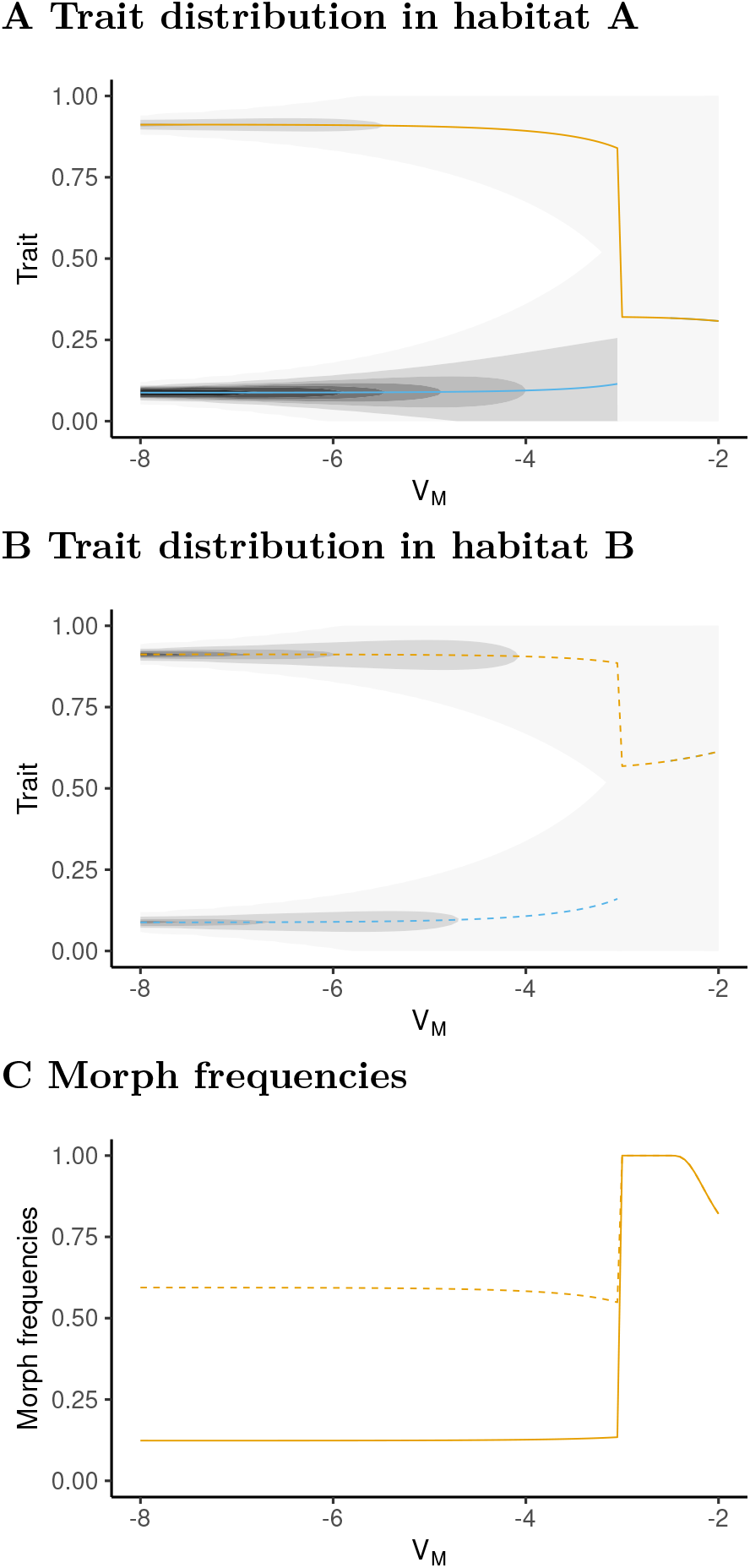
Effect of mutation. The density distribution in habitats A (panel (a)) and B (panel (b)), and the frequencies of morph 2 in each habitat (panel (c); A: solid, B: dashed) are shown as a function of the mutational variance *V_M_* (log scale). When *V_M_* increases, the variance of the distributions increases. There is a threshold at *V_M_* ≈ 10^-3^ above which the oligomorphic approximation breaks down.

Multiplying by 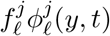 and summing over *ℓ* yields

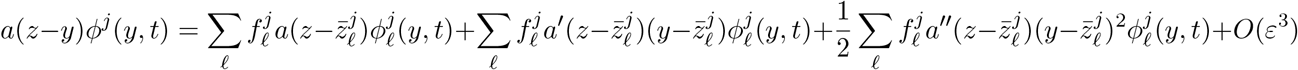

Integrating over *y*, we obtain:

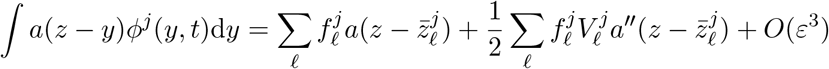

We can then write the vital rates as

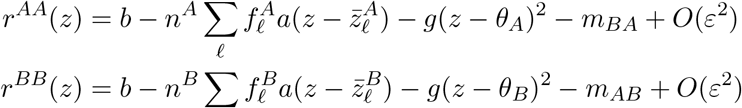

and the partial derivatives:

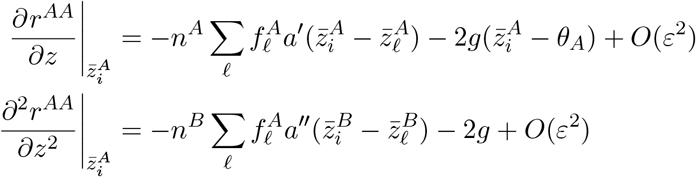

with similar expressions for the partial derivatives in habitat *B*. We can then use these expressions in equations (S.18) to obtain the general oligomorphic approximation of the resource competition model. This is how the numerical simulations in figure 5 in the main text (plain and dashed lines) were performed.

#### Projection on RV space

We can also plug these expressions into equations (S.22) and (S.31) to obtain the projection on RV space:

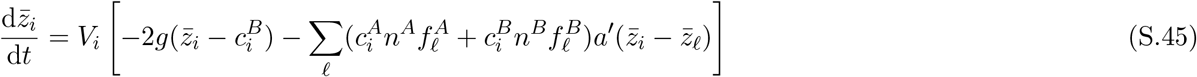

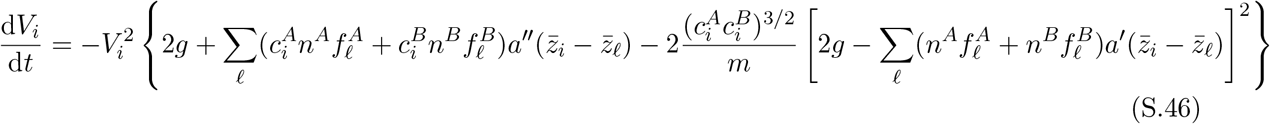

With only one morph and a Gaussian kernel (i.e. *a*(*x*) = *κ* exp(−*x*^2^/(2*ω*^2^))) such that *a*′(0) = 0 and *a*″(0) = −*κ*/*ω*^2^), this yields

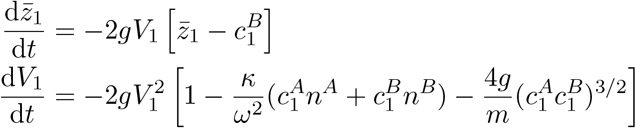

with 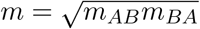,

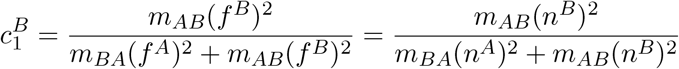

(this directly follows from equation (D.3)) and

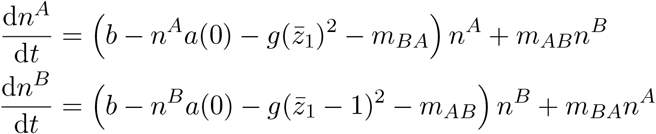

With two morphs and a Gaussian kernel, we have

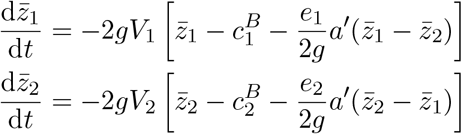

with

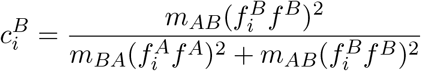

and

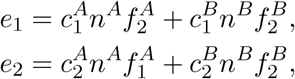

together with the dynamics of morph variances (equation (S.46)) and of *n^k^*(*t*), 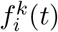 (equations (S.18a)–(S.18c) with 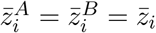). The dotted lines in figure 5 show the results of the numerical integration of this two-morph system.

### S.8 Example 3: A two-habitat resource-consumer model

We now consider a resource-consumer model where the fitness function of individuals with trait *z* in habitat *k* are given by

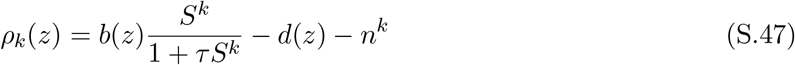

where *S^k^* is the density of resource in habitat *k*, *b*(*z*) is the fecundity rate, *d*(*z*) the mortality rate, and *τ* the handling time (that is we assume a type-II functional response). We assume a trade-off between fecundity and survival, and in particular, for our simulations, we will use

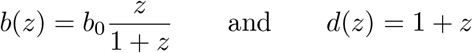

We have the following transition rates

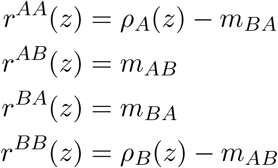

and the eco-evolutionary dynamics of the consumer population can be approximated using equations (S.18), together with the following equations for the dynamics of the resources:

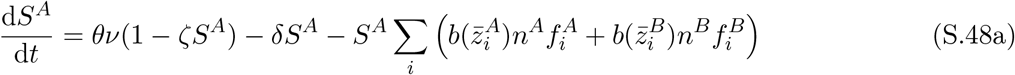

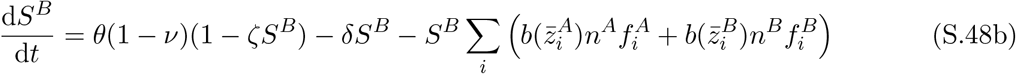

We therefore assume that the resource is produced in each habitat in a biased manner (first term on the right-hand sides), and decays at rate *δ* in both habitats (second term). The third term describes the depletion of resource due to the exploitation by the consumer, and is simply derived by a Taylor-expansion of

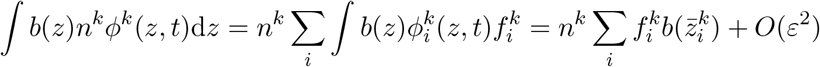

For this example, we are interested in the dynamics of the mean trait in a given habitat, that is across the different peaks of the multi-modal distribution. The mean trait in habitat *A* can be calculated as

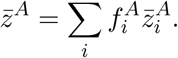

Differentiating this equation yields

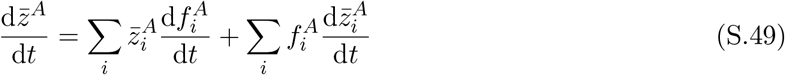

The first term tells us how the mean trait in habitat A changes when the height of the peaks change (e.g. the frequencies). This describes fast dynamics. The second term tells us how the mean trait in habitat A changes when the positions of the peaks change (e.g. the morph means). This describes slow dynamics. To simplify the second term, we use the projection on RV space and write

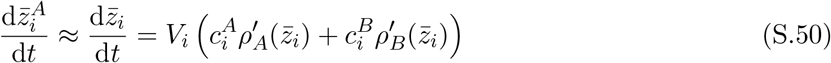

To simplify the first term we assume that we have only two morphs. Then 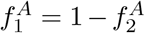 and we obtain

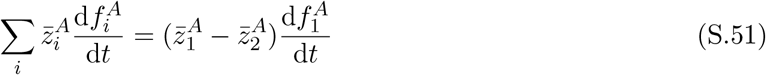

and use equation (S.18b) to obtain

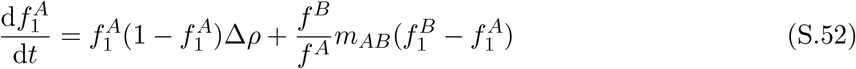

where 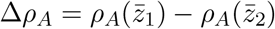 is the average difference in growth rates between the two morphs.

Using the approximation 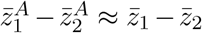 and inserting equations (S.50) and (S.52) into equation (S.49) yields

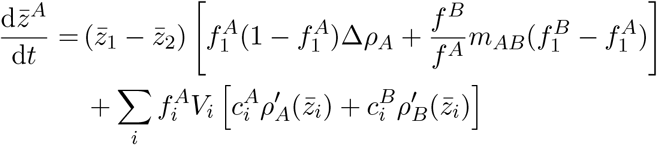

which is equation (37) in the main text.

An interesting limit is when the population eventually becomes monomorphic (say morph 1 goes to fixation), in which case 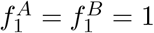 and therefore the first line vanishes, and the dynamics of 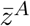 collapse to:

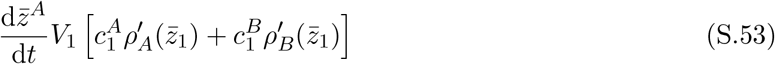

and we have 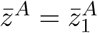. The term between brackets is the selection gradient one would obtain from an invasion analysis in a monomorphic population and allows us to calculate evolutionary singularities. In this model, with the trade-off functions we choose, we obtain the following implicit relationship:

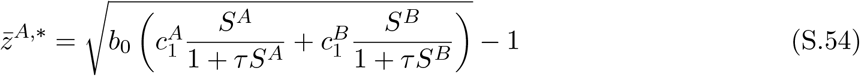

#### Additional figures

In the main text, we discuss a specific scenario and we provide here two additional figures. Figure S.7 presents the same simulation results as in figure 6 in the main text but presents the dynamics of the morph frequencies and morph means, instead of the class-level means and variances. Figure S.8 presents the same scenario as in figure 6 in the main text, but starting from two morphs that have very similar trait values, so that the overall standing variation in the population is small.

#### Parameter values

For this scenario, we use the following parameter values: *θ* = 8, *ζ* = 0.25, *ν* = 0.2, *d* = 1, *δ* = 0.8, *b*_0_ = 7, *τ* = 0.5, *V_M_* = 10^-5^, *m_AB_* = 0.2, *m_BA_* = 0.4. See the companion notebook for more details on the initial conditions.

**Figure S.7:**
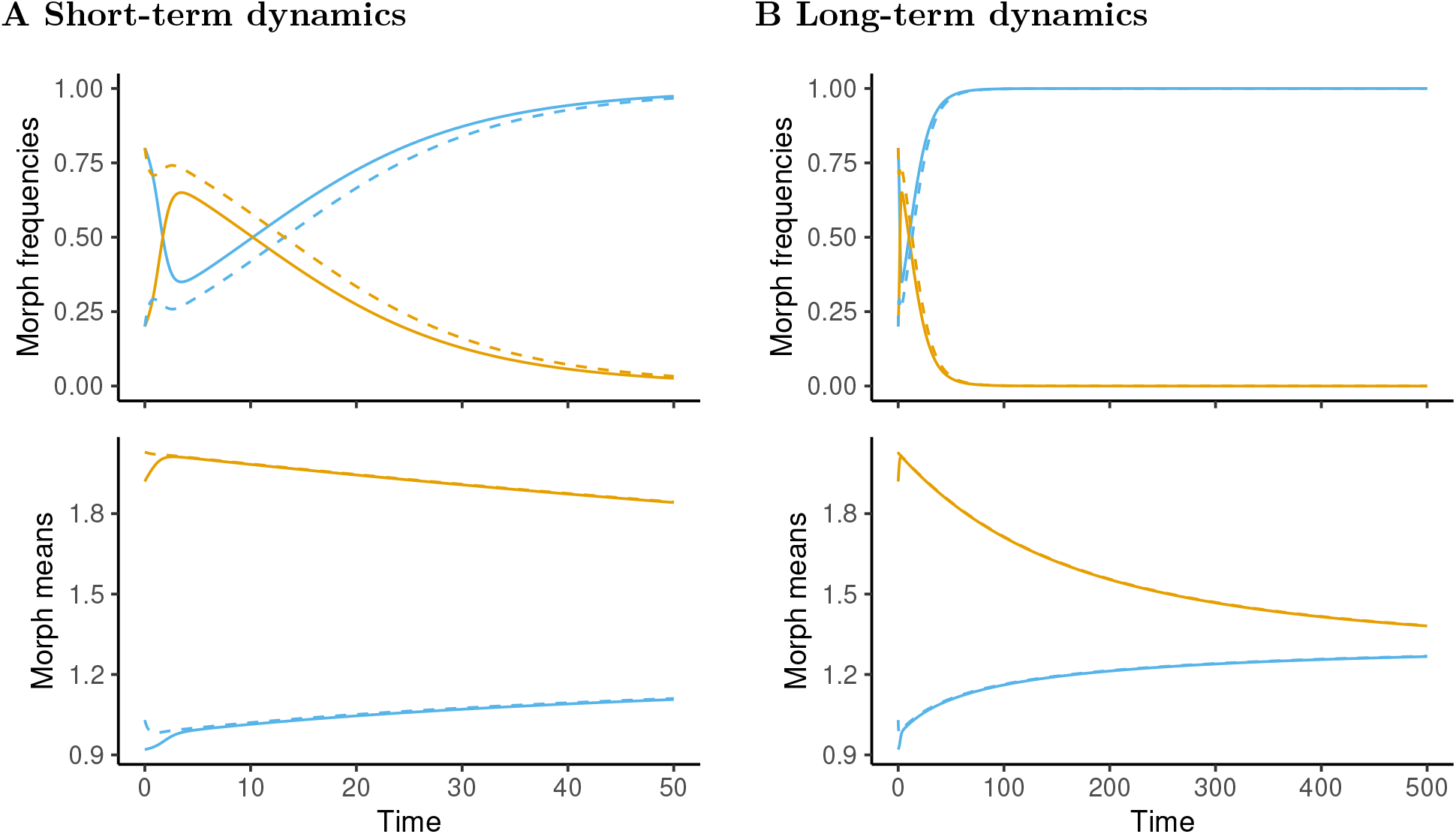
Dynamics of morph frequencies (top panel) and morph means (bottom panel) in the simulations of figure 6 in the main text, for both short-term (a) and long-term (b) dynamics.

**Figure S.8:**
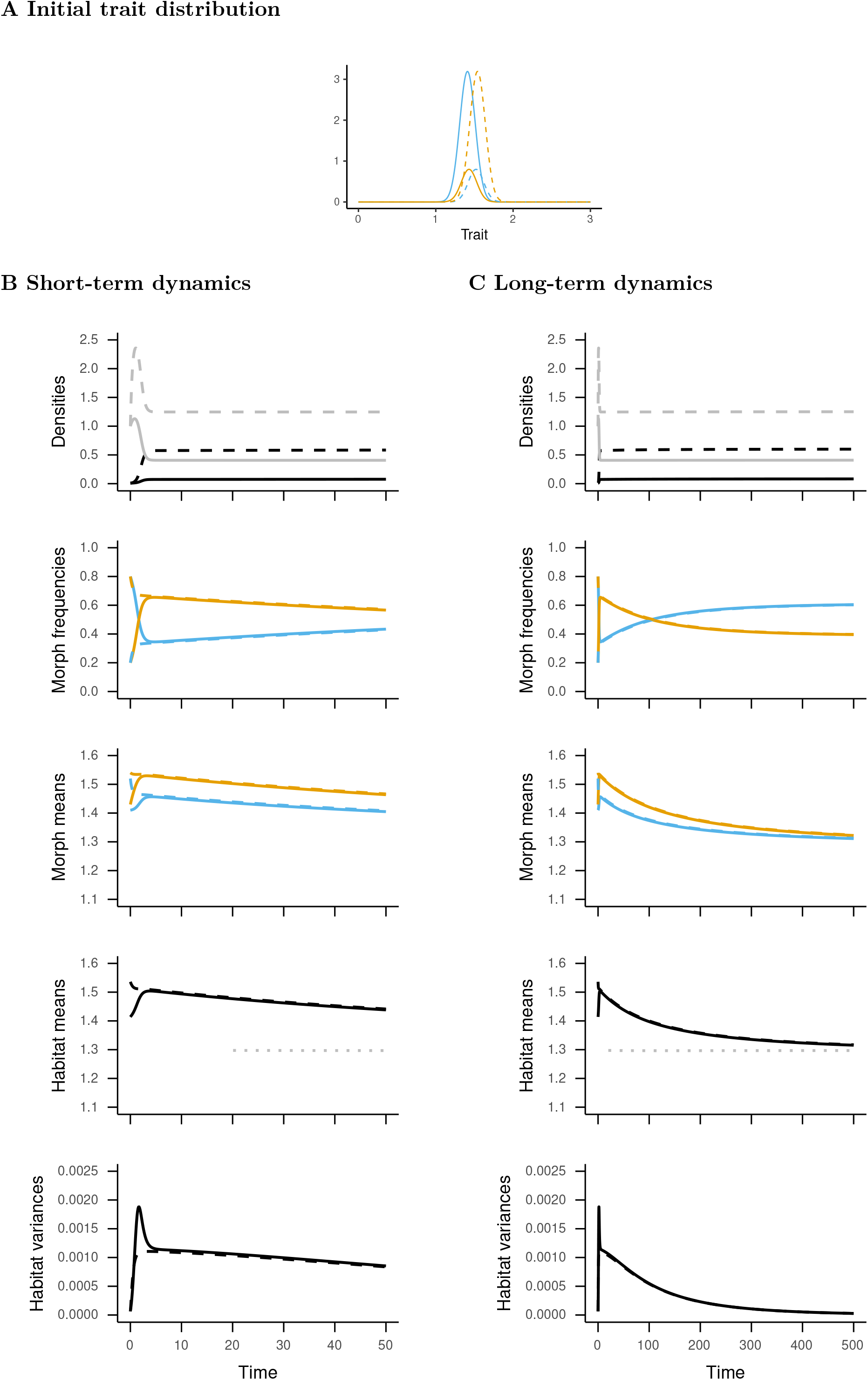
Same as in figure 6 in the main text, but when the two morphs are very close.

